# RelA-SpoT Homologue toxins pyrophosphorylate the CCA end of tRNA to inhibit protein synthesis

**DOI:** 10.1101/2021.04.30.441581

**Authors:** Tatsuaki Kurata, Tetiana Brodiazhenko, Sofia Raquel Alves Oliveira, Mohammad Roghanian, Kathryn Jane Turnbull, Ondřej Bulvas, Hiraku Takada, Hedvig Taman, Andres Ainelo, Radek Pohl, Dominik Rejman, Tanel Tenson, Abel Garcia-Pino, Gemma C. Atkinson, Vasili Hauryliuk

## Abstract

RelA-SpoT Homolog (RSH) enzymes control bacterial physiology through synthesis and degradation of the nucleotide alarmone (p)ppGpp. We recently discovered multiple families of Small Alarmone Synthetase (SAS) RSH acting as toxins of toxin-antitoxin (TA) modules, with the FaRel subfamily of toxSAS abrogating bacterial growth by producing an analogue of (p)ppGpp, (pp)pApp. Here we probe the mechanism of growth arrest employed by four experimentally unexplored subfamilies of toxSAS: FaRel2, PhRel, PhRel2 and CapRel. Surprisingly, all these toxins specifically inhibit protein synthesis. To do so, they transfer a pyrophosphate moiety from ATP to the tRNA 3′ CCA. The modification inhibits both tRNA aminoacylation and the sensing of cellular amino acid starvation by the ribosome-associated RSH RelA. Conversely, we show that some Small Alarmone Hydrolase (SAH) RSH enzymes can reverse the pyrophosphorylation of tRNA to counter the growth inhibition by toxSAS. Collectively, we establish RSHs as a novel class of RNA-modifying enzymes.

## Introduction

Toxin–antitoxin (TA) systems are a class of highly diverse and widespread small operons found in bacterial, archaeal and bacteriophage genomes (Fraikin et al., 2020; Harms et al., 2018). TAs have a broad range of functions, including bacterial defence against bacteriophages, phage competition for infection of bacteria, stabilization of transposons, plasmids and bacterial genomes – all of which rely on the highly potent toxicity of the protein toxin controlled by the protein- or RNA-based antitoxin (Blower et al., 2009; Fiedoruk et al., 2015; Jaffe et al., 1985; Lima-Mendez et al., 2020; Song and Wood, 2020). Many TA toxins are closely evolutionary related to housekeeping enzymes, suggesting a ‘breakaway’ evolutionary path on which a generally harmless enzyme evolves a toxic function that requires tight control by the antitoxin (Burckhardt and Escalante-Semerena, 2020; Garcia-Pino et al., 2014; Jimmy et al., 2020; Koonin and Makarova, 2019; Senissar et al., 2017).

The RelA-SpoT Homolog (RSH) protein family of housekeeping stress-response enzymes was only recently recognised to also contain TA toxins (Jimmy et al., 2020). The function of housekeeping RSHs is to control the cellular levels of alarmone nucleotides ppGpp (guanosine-3’,5’-bisdiphosphate) and pppGpp (guanosine-5’-triposphate-3’-diphosphate) – collectively referred to as (p)ppGpp – with the alarmones, in turn, regulating metabolism, virulence and growth rate, as well as playing an important role in antibiotic and stress tolerance (Gaca et al., 2015; Hauryliuk et al., 2015; Irving et al., 2020; Liu et al., 2015; Zhu et al., 2019). RSH family members can both synthesise (p)ppGpp by transferring the pyrophosphate group of ATP to the 3’ position of either GDP or GTP, and convert it back to GDP/GTP through removal of the 3’ pyrophosphate (Atkinson et al., 2011; Cashel and Gallant, 1969). RSHs can be classified into long multi-domain RSHs – with the archetypical representatives being *Escherichia coli* enzymes RelA (Haseltine and Block, 1973) and SpoT (Xiao et al., 1991) – and short single-domain RSHs (Atkinson et al., 2011). The latter are highly diverse, and can be subdivided into Small Alarmone Synthetases (SAS; 30 distinct subfamilies, with numerous experimentally representatives extensively characterised, including *Staphylococcus aureus* RelP (Geiger et al., 2014; Manav et al., 2018) and *Bacillus subtilis* RelQ (Nanamiya et al., 2008; Steinchen et al., 2015)) and Small Alarmone Hydrolases (SAH; 11 subfamilies) (Jimmy et al., 2020).

Several recent discoveries have sparked interest in how non-(p)ppGpp RSH-mediated chemical catalysis can be weaponised by bacteria for potent growth inhibition. Highly toxic SAS RSH enzymes that are injected by secretion systems (Ahmad et al., 2019) or constitute the toxic components (toxSAS), of toxin-antitoxin (TA) modules (Jimmy et al., 2020) were found to produce a structural analogue of (p)ppGpp – pApp, ppApp and pppApp – collectively constituting (pp)pApp (**Figure S1**). By abrogating *de novo* purine synthesis through orthosteric inhibition of PurF (Ahmad et al., 2019), (pp)pApp inhibits translation, transcription and replication (Jimmy et al., 2020). SAHs have also been shown to catalyse unexpected new reactions. While no (p)ppGpp synthetases are encoded in mammalian genomes, human MESH1 was identified in 2010 as an efficient (p)ppGpp hydrolase (Sun et al., 2010). A decade later, compelling evidence was presented that human MESH1 is a NADPH phosphatase (Ding et al., 2020). Combined with the dramatic evolutionary diversity of this largely experimentally unexplored protein family (Atkinson et al., 2011; Jimmy et al., 2020), these discoveries demonstrate that RSH-mediated catalysis is versatile and that the biological functions of RSH enzymes are clearly not limited to (p)ppGpp metabolism.

We have recently experimentally validated representatives of five toxSAS subfamilies as *bona fide* TA effectors: *Cellulomonas marina* FaRel, *Bacillus subtilis* la1a PhRel2, *Coprobacillus* sp. D7 FaRel2, *Mycobacterium* phage Phrann PhRel (Gp29) and *Mycobacterium tuberculosis* AB308 CapRel (Jimmy et al., 2020). Out of these, only the (pp)pApp-producing *C. marina* FaRel was previously functionally characterised. In this study we uncover surprising non-alarmone chemistry catalysed by previously unexplored FaRel2, PhRel2, PhRel and CapRel enzymes as well as shed light on how the toxicity of non-alarmone toxSASs can be counteracted through the hydrolytic activity of SAHs.

## Results

### Representatives of FaRel2, PhRel, PhRel2 and CapRel toxSAS subfamilies specifically inhibit protein synthesis

While the mechanism of toxicity employed by PhRel, PhRel2, FaRel2 and CapRel toxSAS subfamilies is as yet uncharacterised, we initially assumed that these toxSASs – just as *C. marina* FaRel TA toxin (Jimmy et al., 2020) and the Tas1 toxic RSH effector of *Pseudomonas aeruginosa* Type VI secretion system (Ahmad et al., 2019) – inhibit bacterial growth by producing (pp)pApp. Surprisingly, when we analysed the nucleotide pools of growth-arrested *E. coli* expressing *B. subtilis* la1a PhRel2, we detected no accumulation of (pp)pApp (**Figure 1A** and **Figure S1D**). At the same time, we robustly detected (pp)pApp upon expression of FaRel (**Figure S1E**; the synthesis of (pp)pApp standards is described on **Figure S2**). Similarly, we did not detect (pp)pApp upon expression of *Coprobacillus* sp. D7 FaRel2 either (**Figure S1H**). These results suggested that toxSASs might not universally act via production of (pp)pApp, and therefore, multiple toxSAS subfamilies could have a mechanism of toxicity distinct from that of FaRel and Tas1.

**Figure 1.**
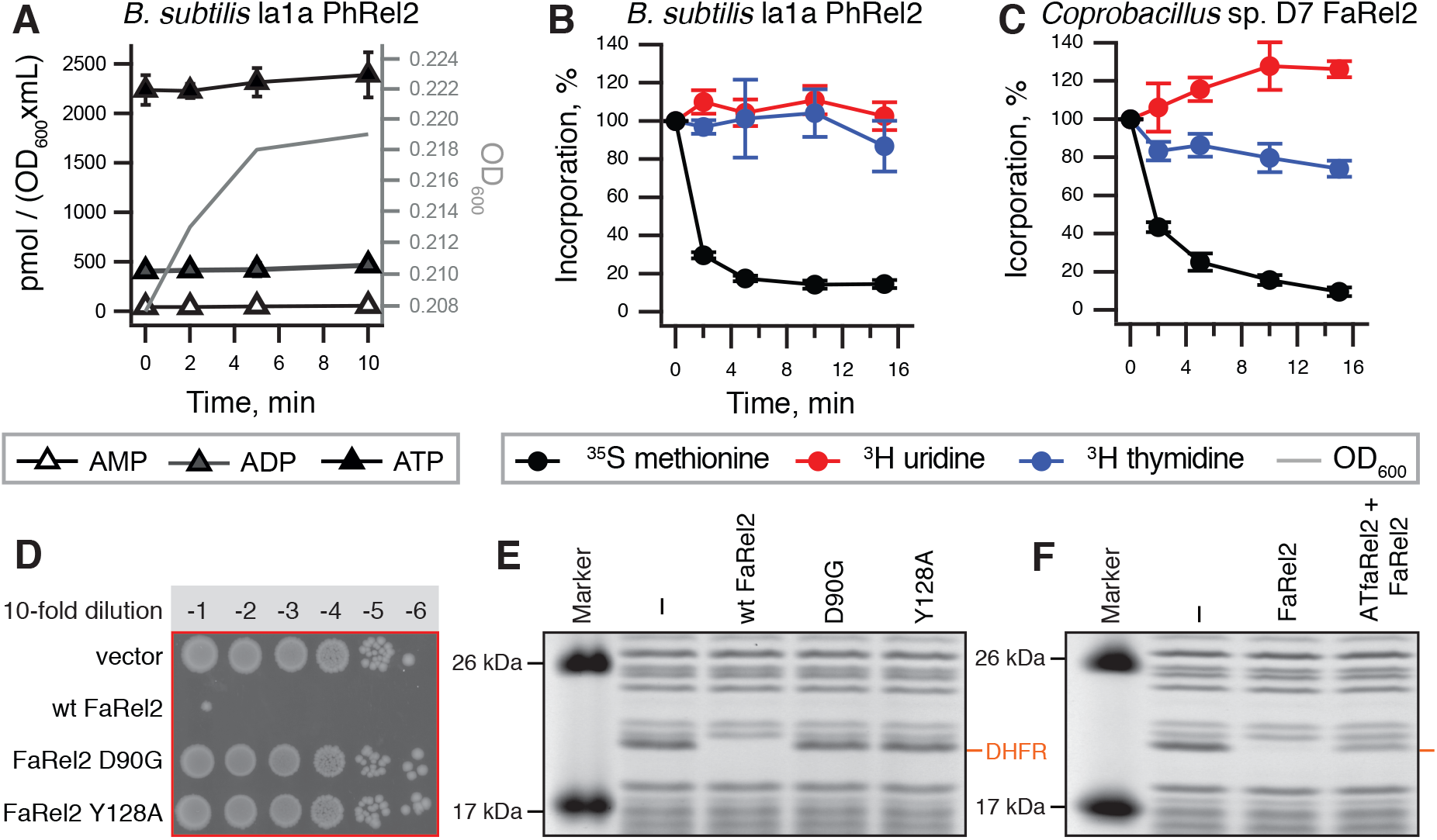
Inhibition of protein synthesis is an evolutionary widespread mechanism of toxSAS-mediated growth arrest. (**A**) The expression of *B. subtilis* la1a PhRel2 does not perturb the adenosine nucleotide pools and (pp)pApp is not detectable upon expression of the toxin. Analogous experiments with *C. marina* FaRel and *Coprobacillus* sp. D7 FaRel2 are presented in **Figure S1**. (**B, C**) Pulse-labelling assays following incorporation of ^35^S-methionine (black traces), ^3^H-uridine (red traces) and ^3^H-thymidine (blue traces). The expression of *B. subtilis* la1a PhRel2 (**B**) and *Coprobacillus* sp. D7 FaRel2 (**C**) from the pBAD33-based constructs was induced with 0.2% L-arabinose. Analogous experiments with *P. aeruginosa* Tas1, *Mycobacterium* phage Phrann PhRel (Gp29) and *M. tuberculosis* AB308 CapRel toxSAS are presented in **Figure S3**. (**D**) D90G and Y128A substitutions render FaRel2 non-toxic. (**E, F**) Cell-free expression assays. Wild-type but not D90G or Y128A substituted FaRel2 abrogates production of DHFR (**E**). The addition of the ATfaRel2 antitoxin counteracts the inhibitory effect of FaRel2 (**F**).

We used metabolic labelling to uncover the effects of as yet uncharacterised toxSASs on translation (by following incorporation of ^35^S-methionine in proteins), transcription (incorporation of ^3^H-uridine in RNA) and replication (incorporation of ^3^H-thymidine in DNA). In stark contrast to FaRel (Jimmy et al., 2020) and Tas1 (Ahmad et al., 2019) which both inhibit all the three processes (**Figure S3A**), representatives of all four unexplored toxSAS subfamilies specifically inhibited translation. The strongest inhibition was observed for *B. subtilis* la1a PhRel2 and *Coprobacillus* sp. D7 FaRel2 (**Figure 1C**,**B**), while *M. tuberculosis* AB308 CapRel and *Mycobacterium* phage Phrann superinfection immunity protein PhRel (Gp29) (Dedrick et al., 2017) had a weaker, but still specific effect on protein synthesis (**Figure S3B**,**C**). Interestingly, upon induction of FaRel2, ^3^H-uridine incorporation increased. This is likely due to abrogation of ATP consumption upon cessation of translation, resulting in increased transcription; we earlier observed a similar effect upon specific inhibition of translation by the antibiotic kanamycin (Jimmy et al., 2020). Collectively, our results suggested that specific inhibition of translation is a common mode of toxSAS toxicity, with the FaRel toxSAS subfamily deviating from this common *modus operandi*.

We next tested whether inhibition of translation by toxSASs is mediated by a direct mechanism using production of dihydrofolate reductase, DHFR, in a reconstituted cell-free protein synthesis system from *E. coli* components (PURE) (Shimizu et al., 2001) as a readout of toxSAS activity. Although purification of toxSAS enzymes is exceedingly challenging (Jimmy et al., 2020), we succeeded in purifying enzymatically-competent C-terminally FLAG_3_-tagged *Coprobacillus* sp. D7 FaRel2 toxin through α-FLAG_3_-immunoprecipitation (**Figure S4**). As we have shown earlier, the FLAG_3_-tag does not interfere with toxicity of FaRel2 or the ability of the antitoxin to counteract the protein (Jimmy et al., 2020). As a specificity control, we used catalytically compromised FaRel2 variants Y128A (predicted to disrupt the stacking interaction with the pyrophosphate acceptor substrate (Steinchen et al., 2018)) and D90G (predicted to compromise the coordination of the Mg^2+^ ion (Steinchen et al., 2015)). Both of the substituted residues are highly conserved amongst SAS RSH enzymes (**Figure S5A**) and mutant variants are non-toxic when expressed in *E. coli* (**Figure 1D**). The addition of wild-type – but not D90G or Y128A – FaRel2 to the PURE system abrogated DHFR production (**Figure 1E, Figure S6A**). The addition of the ATfaRel2 antitoxin which acts though sequestering FaRel2 into inactive complex (Jimmy et al., 2020) counteracted the inhibitory effect of FaRel2 (**Figure 1F**). We concluded that this toxSAS, indeed, directly targets the protein synthesis machinery.

### *Coprobacillus* sp. D7 FaRel2 specifically modifies the tRNA 3′ CCA end to abrogate aminoacylation

Inhibition of protein production is a common means of toxicity in TA systems, with the toxic components often acting via modification of tRNA, such as cleavage (employed by VapC toxins (Cruz et al., 2015; Winther and Gerdes, 2011)), acetylation of the attached amino acid (as seen with GNAT toxins (Cheverton et al., 2016; Jurenas et al., 2017)) or inactivation of the 3′ CCA end through the addition of pyrimidines (employed by MenT_3_ (Cai et al., 2020)). RSH enzymes have never previously been shown to catalyse synthesis of any other products than hyperphosphorylated nucleotides (pp)pGpp and (pp)pApp. However, one could imagine that the pyrophosphate group of the ATP donor could be transferred onto the ribose position of the 3′ terminal adenosine of tRNA instead of the corresponding 3′ ribose position of the ATP/ADP substrate used by Tas1/FaRel to produce (pp)pApp. Since the availability of this 3′ hydroxyl group is essential for tRNA aminoacylation, the modification would efficiently inhibit protein synthesis.

We tested this hypothesis using deacylated *E. coli* tRNA as a substrate and γ-^32^P ATP as a donor of radioactively labelled pyrophosphate moiety. In the presence of γ-^32^P ATP FaRel2 efficiently radiolabels both initiator tRNA_i_ ^fMet^ (**Figure 2A**,**B**) and elongator tRNA^Phe^ (**Figure 2B**). The tRNA-labelling activity is lost in D90G and Y128A FaRel variants (**Figure 2C**), and is specifically counteracted by both the AtFaRel2 Type II antitoxin and tRNA aminoacylation (**Figure 2D**,**E**). The latter result strongly suggests that the ^32^P label is transferred by FaRel2 onto the 3′ hydroxyl group of the tRNA terminal adenine residue that acts as an amino acid acceptor. To probe this experimentally, we tested the effect of tRNA modification by FaRel2 on aminoacylation of tRNA^Phe^. The aminoacylation reaction was readily abrogated by FaRel2 (**Figure 2F**) in a strictly ATP-dependent manner (**Figure 2G**), thus explaining the molecular mechanism of translational arrest by this toxSAS. In principle, the FaRel2-modified phospho-tRNA might not just be incompetent in aminoacylation, but also actively toxic to translation due to, for instance, stable binding to the ribosomal A-site or elongation factor EF-Tu. To test this hypothesis, we first titrated total tRNA in the PURE system and identified the near-saturating tRNA concentration (50 µM) at which, we reasoned, the system would be most sensitive to inhibition (**Figure S6B**). However, added in concentrations up to 14 µM, phospho-tRNA^Phe^ had no effect on DHFR synthesis (**Figure S6C**), suggesting that while phospho-tRNA is translation-incompetent, it is not inhibitory to the protein synthesis machinery. Rather, the toxicity likely results from a depletion of chargeable tRNA in the cell.

**Figure 2.**
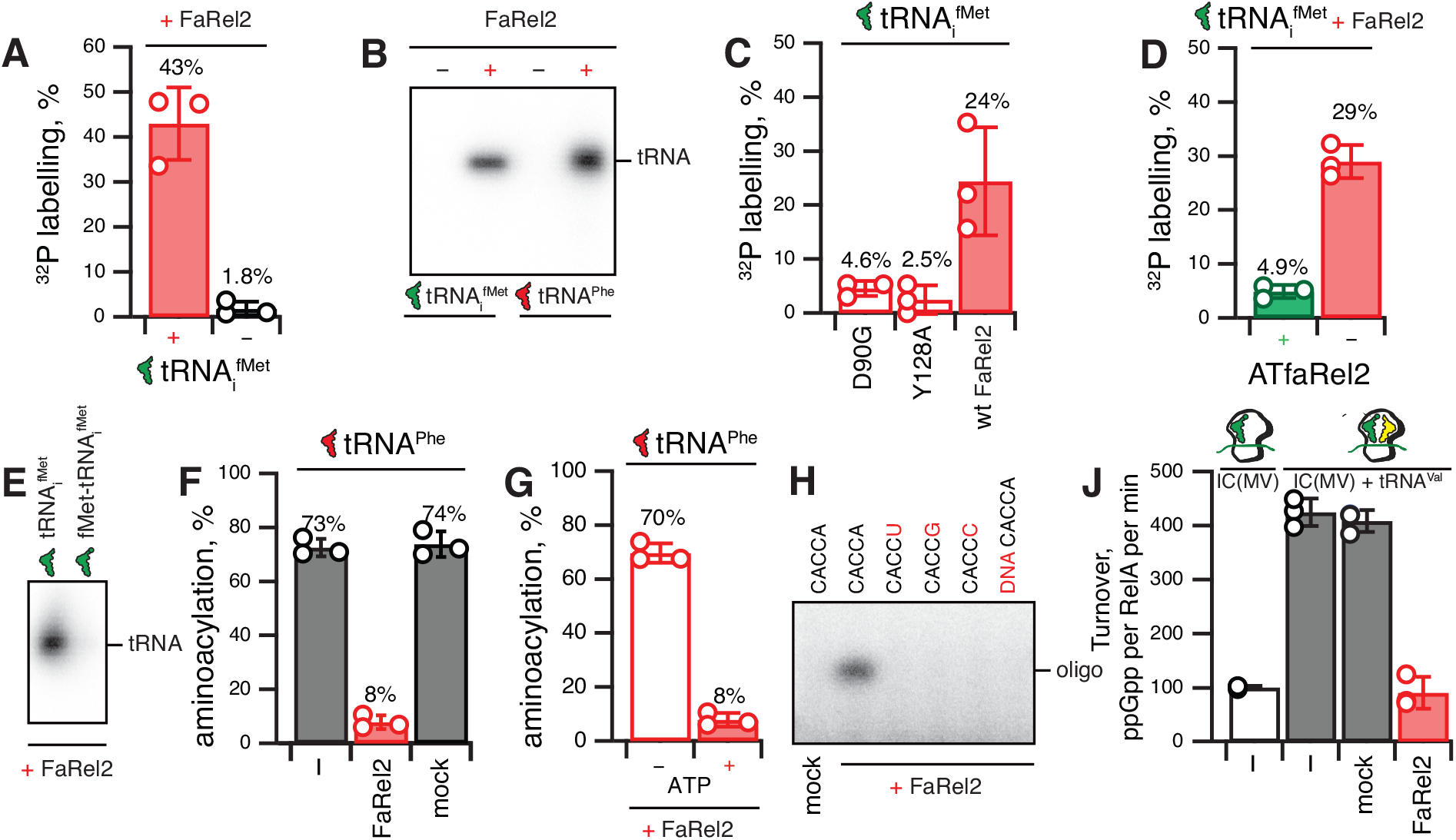
The FaRel2 toxSAS toxin modifies the 3′ CCA end of tRNA to inhibit aminoacylation and the sensing of amino acid starvation by RelA. (**A, B**) A reconstituted ^32^P transfer reaction using FaRel2 and either tRNA_i_^fMet^ or tRNA^Phe^ as a substrate. (**C**) Non-toxic D90G and Y128A FaRel2 mutants are compromised in their ability to modify tRNA. (**D**) The ATfaRel2 antitoxin counteracts tRNA modification by FaRel2. (**E**) Acylated fMet-tRNA_i_^fMet^ is refractory to modification by FaRel2. (**F**) tRNA^Phe^ modification by FaRel2 inhibits aminoacylation. As specificity controls, the reactions were supplemented either with mock protein preparation from *E. coli* strain transformed with an empty plasmid vector (mock) or HEPES:Polymix buffer (–). (**G**) Inhibition of tRNA^Phe^ aminoacylation by FaRel2 is strictly ATP-dependent. (**H**) 3′ adenosine defines the specificity of modification by FaRel2, as tested using a set of model 5′-CACCN-3′ RNA oligonucleotides. 5′-CACCA-3′ DNA does not serve as a substrate for FaRel2. (**J**) FaRel2 abrogates the stimulatory effect of deacylated tRNA^Val^ on ppGpp synthesis by *E. coli* RelA in the presence of 70S ribosomal initiation complexes. Error bars represent standard deviations of the mean. The mock sample was produced by immunoprecipitation using *E. coli* cells transformed with a plasmid vector not expressing FLAG_3_-tagged FaRel2.

To test the generality of toxSAS-mediated translation inhibition via tRNA CCA pyrophosphorylation, we purified and tested *B. subtilis* la1a PhRel2. Similarly to *Coprobacillus* FaRel2, PhRel2 efficiently abrogates protein synthesis in the PURE system (**Figure S6D**) and γ-^32^P-labels both tRNA_i_ ^fMet^ and tRNA^Phe^ (**Figure S6E**).

Finally, we assessed the specificity of tRNA modification by FaRel2. Both deacylated initiator tRNA_i_^fMet^ and elongator tRNA^Phe^ were labelled with ^32^P by FaRel2 (**Figure 2B**), which suggests that the 3′ CCA could be sufficient for recognition of deacylated tRNA. To test this hypothesis and probe the specificity of the toxSAS for tRNA’s 3′ adenine residue we performed radiolabelling experiments with a set of synthetic 5′-CACCN-3′ RNA pentanucleotides containing both the 3′ A as well as 3′ C, G and U (**Figure 2H**). Only one of the four RNA substrates, CACCA, was labelled with ^32^P, suggesting specificity for 3′ adenine. At the same time, the CACCA DNA oligonucleotide did not serve as a FaRel2 substrate, suggesting a functional importance of the 2′ hydroxyl group of the 3′ adenine (**Figure 2H**). This result is in good agreement with a recent report demonstrating that (p)ppGpp-synthesising RSH enzymes cannot catalyse the transfer of the pyrophosphate group of the ATP donor to dGTP instead of the GTP substrate (Patil et al., 2020).

### tRNA 3′ CCA end modification by FaRel2 abrogates ribosome-dependent activation of (p)ppGpp synthesis by amino acid starvation sensor RelA

In Gammaproteobacteria such as *E. coli*, amino acid limitation is sensed by a housekeeping multidomain RSH enzyme RelA (Atkinson et al., 2011). This ribosome-associated factor inspects the aminoacylation status of the 3′ CCA of the A-site tRNA (Arenz et al., 2016; Brown et al., 2016; Loveland et al., 2016), and, upon detecting deacylated tRNA, synthesises the (p)ppGpp alarmone (Haseltine and Block, 1973). While the free 3′ OH moiety of the terminal adenosine residue is essential for full activation of RelA’s synthesis activity by tRNA on the ribosome (Sprinzl and Richter, 1976), RelA is still activated by the 70S ribosome, although to a lesser extent if activation by tRNA is compromised by the antibiotics thiostrepton (Kudrin et al., 2017) and tetracycline (Kudrin et al., 2018).

Using a reconstituted *E. coli* biochemical system (Kudrin et al., 2018) we tested the effect of FaRel2 on RelA activation by deacylated tRNA of starved ribosomal complexes (**Figure 2J**). FaRel2 efficiently abrogated activation of RelA by tRNA, reducing RelA activity to the levels observed in the presence of 70S initiation complexes lacking the A-site deacylated tRNA. Thus, not only does FaRel2 *not* produce an alarmone, it also could prevent the housekeeping RSH cellular machinery from being activated by starved ribosomes to produce the (p)ppGpp alarmone.

### Small Alarmone Synthetase (SAH) RSH enzymes can restore tRNA aminoacylation competence of FaRel2-modified tRNA

When co-expressed with *Coprobacillus* FaRel2, human MESH1 and *C. marina* ATfaRel hydrolysis-only Small Alarmone Synthetase (SAH) enzymes can efficiently counteract the growth inhibition by the toxSAS (**Figure 3A**) (Jimmy et al., 2020). This detoxification activity suggests that MESH1 and ATfaRel can recycle pyrophosphorylated tRNA back to translation-competent deacylated tRNA.

**Figure 3.**
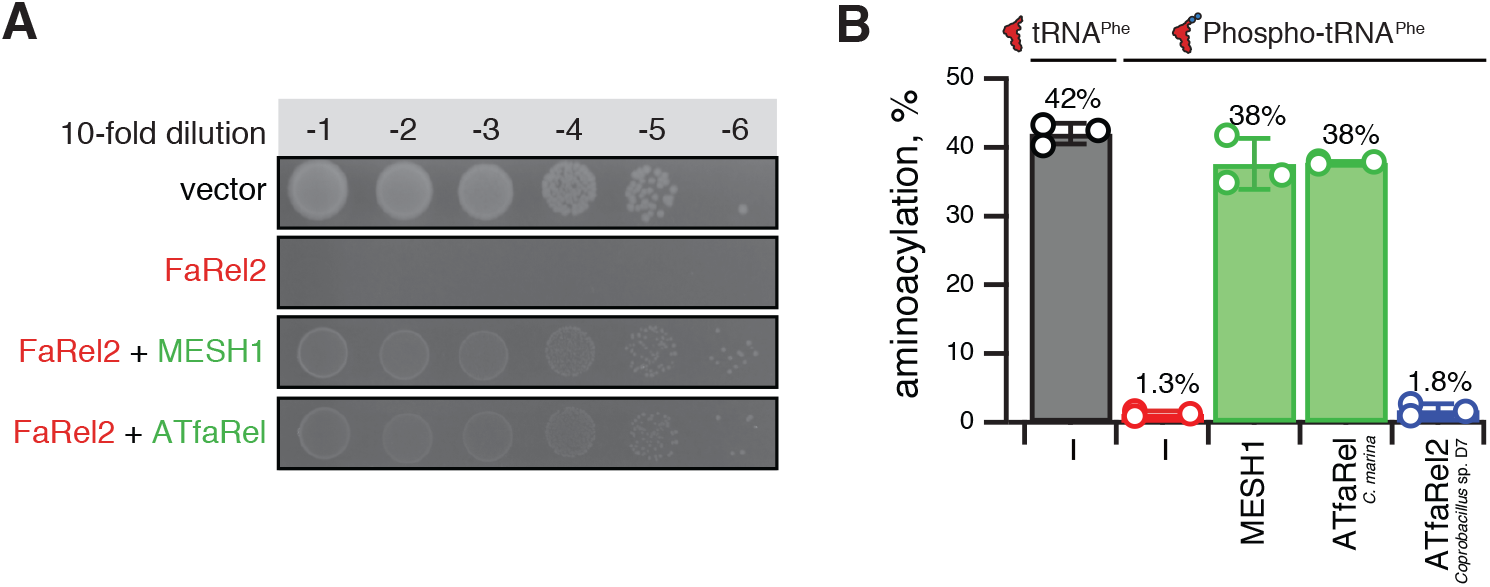
*C. marina* ATfaRel and MESH1 SAH enzymes detoxify FaRel2 through restoration of tRNA aminoacylation. (**A**) Co-expression of human MESH1 and *C. marina* ATfaRel SAH enzymes counteracts the toxicity of *Coprobacillus* sp. D7 FaRel2 toxSAS. The SAH enzymes and FaRel2 toxSAS were induced and expressed from different plasmids, pMG25 and pMR33 derivatives respectively. (**B**) SAH enzymes MESH1 and ATfaRel but not cognate ATfaRel2 Type II antitoxin restore aminoacylation of tRNA^Phe^ abrogated by FaRel2 by CCA pyrophosphorylation. As specificity controls the reactions were supplemented with HEPES:Polymix buffer (–).

To probe this conjecture experimentally, we pyrophosphorylated tRNA^Phe^ *Coprobacillus* FaRel2, isolated the modified tRNA, and tested whether the human MESH1, *C. marina* ATfaRel or cognate Type II *Coprobacillus* AtFaRel2 Type II antitoxin (not an SAH) could restore the aminoacylation activity of pyrophosphorylated tRNA^Phe^ (**Figure 3B**). In excellent agreement with our microbiological results (Jimmy et al., 2020) and consistent with CCA pyrophosphorylation being the cause of growth arrest by FaRel2, both tested SAH enzymes restore the tRNA^Phe^ aminoacylation. At the same time, we detected no effect upon the addition of the ATfaRel2 antitoxin which neutralises FaRel2 by sequestering the toxSAS into an inactive complex (Jimmy et al., 2020), and, therefore, is not expected to restore the aminoacylation competence of FaRel2-modified tRNA.

### Mapping the tRNA 3′ CCA interaction by FaRel2 through molecular docking and mutagenesis

To gain structural insight into the mechanism of tRNA substrate recognition by FaRel2, we used the Rosetta suite (Song et al., 2013) to model the structure of *Coprobacillus* FaRel2 based on the structures of *S. aureus* housekeeping SAS RelP (Manav et al., 2018) and *B. subtilis* SAS RelQ (Steinchen et al., 2015), PDBIDs 6FGK and 6EWZ, respectively. The model predicted by Rosetta was then used to dock deacylated tRNA^Phe^ into the active site as implemented in the HADDOCK suite (van Zundert et al., 2016). As the only distance restraint in the docking experiment we used the necessary proximity of FaRel2 Y128 to the CCA-adenine.

The resulting model of the FaRel2-tRNA complex reveals that a cluster of basic residues accommodate the acceptor stem guiding the CCA end into the active site of the enzyme (**Figure 4A**). In such arrangement, the orientation of the 3′-adenosine is reminiscent of the way GDP is coordinated in the active site of housekeeping SAS *S. aureus* RelP (Manav et al., 2018), next to the binding site of ATP, the pyrophosphate donor. Conversely, the analysis of the electrostatic surface profile of RelP and RelQ shows a charge reversal in the same region (**Figure S5BC**). The presence of a more acidic region in these SASs correlates with a lack of tRNA-pyrophosphorylation activity of these enzymes which would likely be incompatible with tRNA binding.

**Figure 4.**
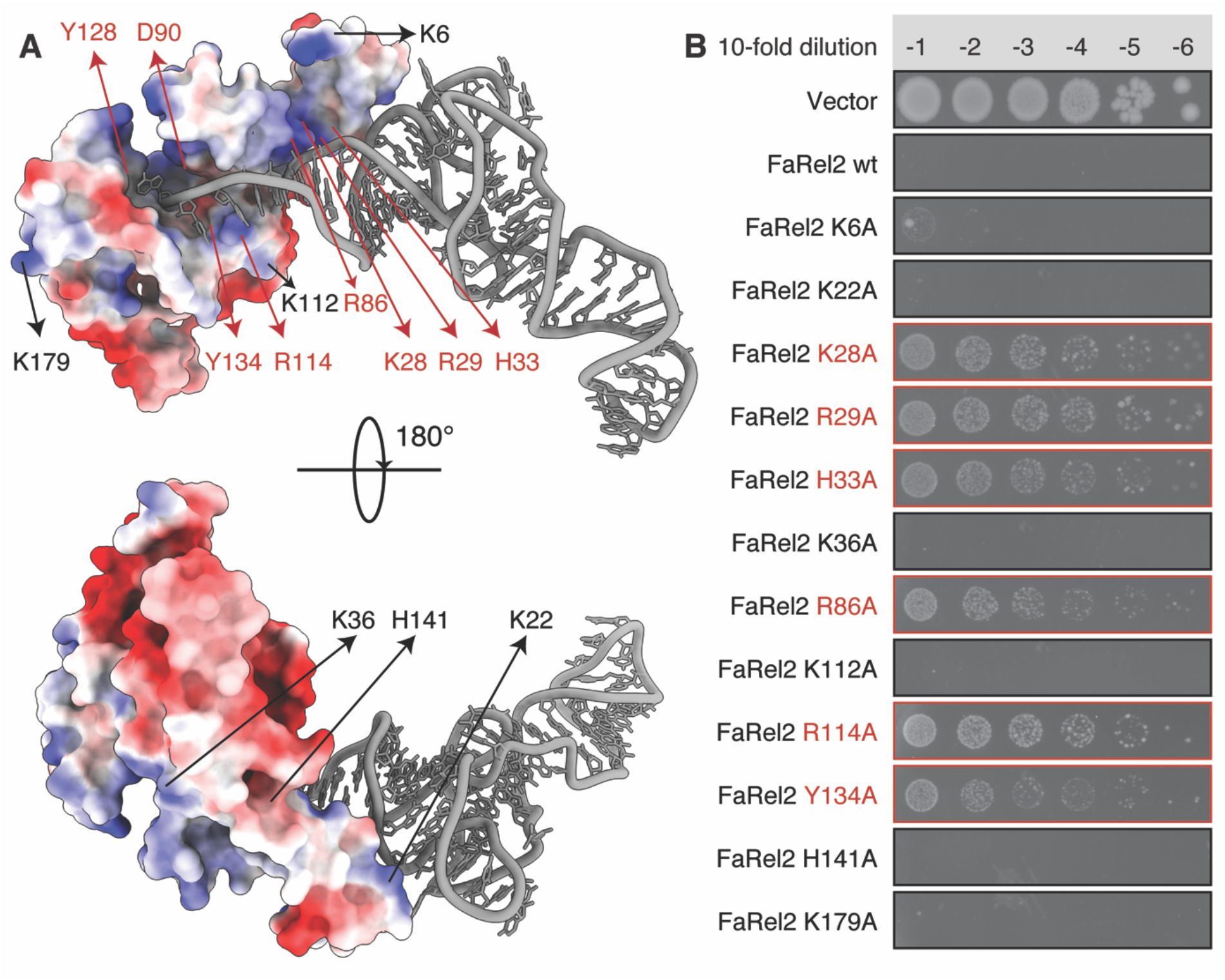
Mutational mapping of the predicted FaRel2: 3′ CCA tRNA interface. (**A**) Surface representation of the model of the FaRel2:tRNA^Phe^ complex. The surface is colored on the basis of electrostatic potential. The phosphodiester backbone of the bound tRNA^Phe^ complements a cluster of positive charges at the active site exit of the enzyme. The predicted FaRel2:tRNA^Phe^ interface involves residues K28, R29, H33, R86, R114 and Y134. While these which guide the CCA end into the active site, functionally essential residue Y128 coordinates the 3′ adenosine of the CCA. (**B**) Ten-fold dilutions of overnight cultures of *E. coli* strains transformed with the pBAD33 vector plasmid or derivatives expressing either wild-type *faRel2* or FaRel2 variants with Ala substitutions at the predicted tRNA-binding interface (K28, R29, H33, R86, R114 and Y134), neighbouring residues (K6, K22, K112 and H141), and positively charged residues outside the binding region (H36 and K179). The latter were served as negative controls. Substitutions at predicted tRNA-binding interface specifically abrogate toxicity of FaRel2.

We probed our molecular model through point mutations in this recognizable basic patch, using FaRel2 toxicity as a readout of intact efficient tRNA recognition (**Figure 4B**). As predicted from our model, Ala-substitutions in the basic cluster located towards the N-terminus of the toxin that includes K28 and K29 abolished the toxicity of FaRel2. Ala-substitutions in the outside rim of the active site (including R114 and Y134) which were predicted to contact the tRNA in our interaction model also compromise the toxicity of FaRel2. These residues are all outside the active site and would not be involved directly in catalysis, thus their effect on toxicity is likely related with tRNA binding. Finally, as a control we confirmed that Ala-substitution of basic residues throughout the surface of FaRel2 had no effect on toxicity (**Figure 4B**).

## Discussion

TA toxins belonging to the same protein family can display relaxed specificity towards their targets (Goeders et al., 2013; Harms et al., 2018; Page and Peti, 2016; Schureck et al., 2015; Yamaguchi and Inouye, 2009) or even enzymatically modify clearly distinct classes of substrates (Burckhardt and Escalante-Semerena, 2020; Castro-Roa et al., 2013; Harms et al., 2015; Jurenaite et al., 2013; Jurenas et al., 2017). A classic example is the GCN5-related N-acetyltransferase (GNAT) TA toxins – a versatile family of enzymes unrelated to RSHs. While GNAT TA toxins inhibit protein synthesis by acetylating aminoacyl-tRNAs (Cheverton et al., 2016; Jurenas et al., 2017), the majority of non-toxic GNATs modify small molecules such as polyamines, antibiotics, phospholipids and amino acids (Burckhardt and Escalante-Semerena, 2020). This substrate specificity spectrum – toxicity mediated via tRNA modification combined with non-toxic modification of small molecule substrates – is strikingly similar to what we present here for RSH enzymes.

Our results uncover a novel enzymatic activity of some toxSAS RSHs to efficiently abrogate translation. We propose the following model of substrate specificity change within the diversity synthetase-competent RSH enzymes (**Figure 5**). The vast majority of RSH synthetases specifically recognise the guanosine residue of the nucleotide (GTP or GDP) substrate to catalyse the synthesis of the housekeeping alarmone (p)ppGpp (**Figure 5A**). In toxic SAS enzymes such *C. marina* FaRel and *P. aeruginosa* Tas1, the substrate specificity is either relaxed, allowing synthesis of both (p)ppGpp and (pp)pApp (toxSAS FaRel (Jimmy et al., 2020)) or switched to specific synthesis of (pp)pApp (Tas1 (Ahmad et al., 2019)) (**Figure 5A**).

**Figure 5.**
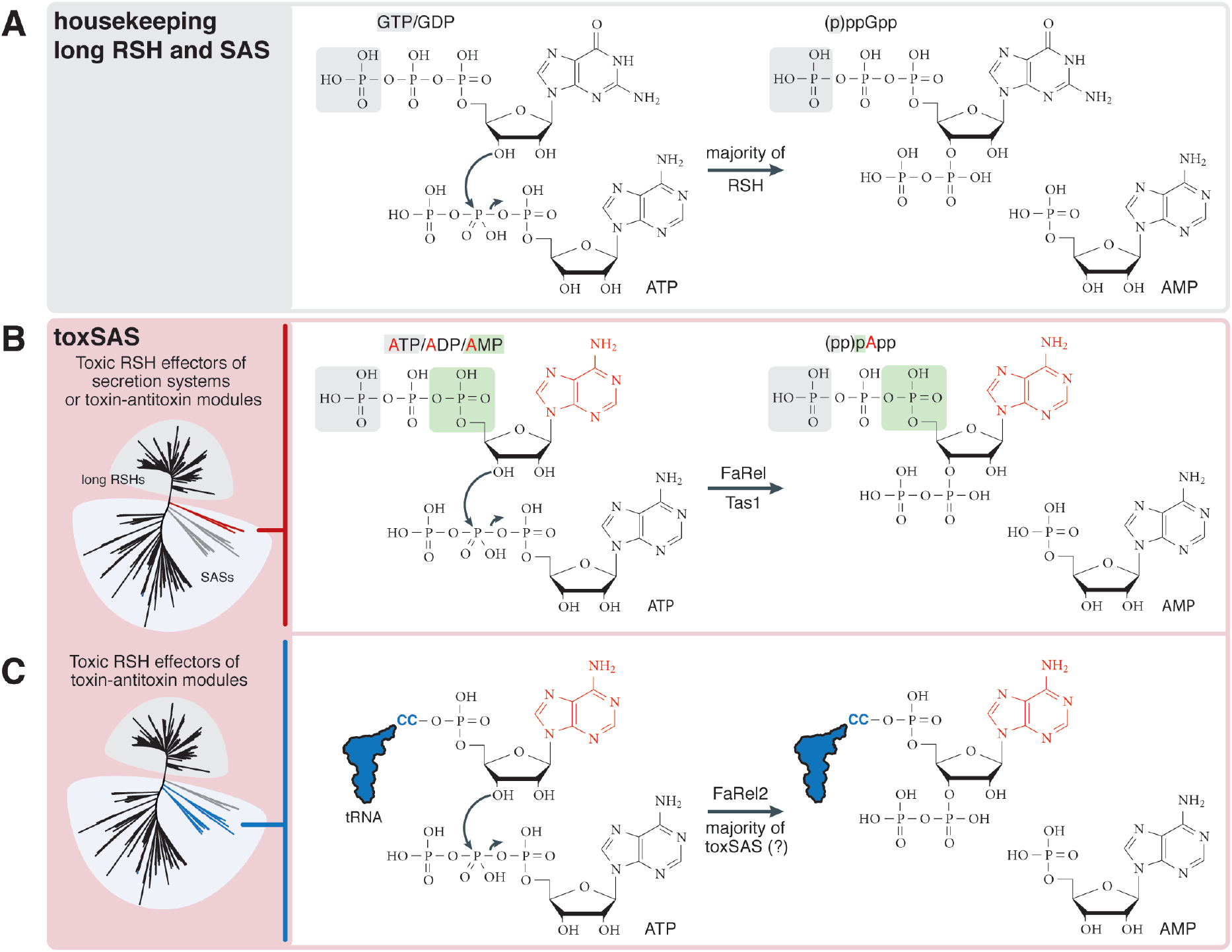
RSH differences in substrate specificity, from nucleotide-mediated signalling via production of (p)ppGpp and (pp)pApp alarmones to toxic modification of the tRNA 3′ CCA end. (**A**) Housekeeping RSHs synthesise (p)ppGpp by transferring the pyrophosphate group of ATP onto the 3′ ribose position of either GDP or GTP, and degrade the alarmone by hydrolysing the nucleotide back to GDP or GTP. (**B**) The substrate specificity of the Tas1 toxic SAS secretion system effectors and FaRel toxic components of toxin-antitoxin systems deviate from strict recognition of the guanine moiety of GDP/GTP employed by ‘housekeeping’ RSHs in favour of the adenine moiety of ATP/ADT/AMP to produce toxic (pp)pApp alarmones. (**C**) The majority of identified toxSAS subfamilies specifically inhibit protein synthesis. FaRel2 toxic components of toxin-antitoxin systems recognise the adenine moiety of tRNA 3′ CCA instead of ATP/ADT/AMP nucleotides, and transfer the pyrophosphate group of ATP onto the 3′ ribose position tRNA 3′ terminal adenosine. This modification abrogates both tRNA aminoacylation and recognition by the amino acid sensor RSH RelA.

Translation-inhibiting toxSASs belong to SAS subfamilies that are found in various major phyla of Gram-positive and -negative bacteria, including Firmicutes, Actinobacteria, Proteobacteria, Bacteroidetes, Acidobacteria, Planctomycetes and Cyanobacteria, as well as multiple bacteriophages and even some archaea (Jimmy et al., 2020). Sequence alignment of the SYNTH domain of toxSASs and other SASs shows that while there are strongly differentially conserved motifs in (pp)pApp-synthesising toxSASs relative to other RSHs, there is – surprisingly – not a clear sequence signature of toxSASs that use tRNA as a substrate (**Figure S5**). Indeed, there is no particular support for monophyly of all toxSASs targeting translation in phylogenetic analysis of RSHs (although there is support for two monophyletic clades comprising FpRel2+PhRel2+FaRel2 and CapRel+PhRel) (Jimmy et al., 2020). The position of the tRNA accepting toxSAS clades at roughly the midpoint of the RSH tree tempts us to speculate that the ancestral function of the SYNTH domain at a time predating the last universal common ancestor (LUCA) could have been pyrophosphorylation of RNA, rather than (pp)pNpp synthesis.

### Perspective and limitations

With 30 distinct SAS subfamilies identified to date (Jimmy et al., 2020), it is likely we are yet to discover the full spectrum of chemical reactions catalysed by the evolutionary versatile RSH synthetase domain. As we show here, pyrophosphorylated tRNA 3′ CCA end can serve as a substrate for SAH enzymes human MESH1 and *C. marina* ATfaRel (**Figure 3B**). This expands the spectrum of known hydrolysis reactions catalysed by RSH beyond hydrolysis of (pyro)phosphorylated nucleotides, indicating a possible new role of RSH hydrolases as RNA-modifying enzymes with a 3′-phosphatase activity similar to that of T4 polynucleotide kinase, Pnk.

This study lays the foundations for the future studies of the non-alarmone chemistry catalysed by bacterial and viral RSH enzymes. Dedicated structural studies are essential for further rationalising our results on the molecular level. All of the experiments presented in the current study are rely on heterologous expression in *E. coli* or utilise reconstituted biochemical system from *E. coli* components. In order to understand the biological roles of toxSAS TAs it will be essential to study these effectors and the antitoxins neutralising them in native bacterial and viral hosts.

## ACKNOWLEDGMENTS

We are grateful to Protein Expertise Platform (PEP) at Umeå University and Mikael Lindberg for constructing plasmids, Andrey Chabes, Nasim Sabouri and Ikenna Obi for providing γ-^32^P ATP, to Steffi Jimmy and Constantine Stavropoulos their contributions at the initial stages of this project. This work was supported by the funds from European Regional Development Fund through the Centre of Excellence for Molecular Cell Technology (VH and TT, 2014-2020.4.01.15-0013); the Molecular Infection Medicine Sweden (MIMS) (VH); The Estonian Research Council (PRG335 to TT and VH); The Swedish Research Council (Vetenskapsrådet; 2017-03783 to VH and 2019-01085 to GCA); The Ragnar Söderberg foundation (VH); MIMS Excellence by Choice Postdoctoral Fellowship Programme (postdoctoral grant 2018 to MR); The Kempe Foundation (SMK-1858.3 to GCA); Carl Tryggers Stiftelse för Vetenskaplig Forskning (19-24 to GCA). The Czech ministry of Education and Sport (grant number 8F19006 to DR) and the Swedish Research Council (2018-00956 to VH) within the RIBOTARGET consortium under the framework of JPIAMR; Umeå Centre for Microbial Research (UCMR) (postdoctoral grant 2017 to HT); the Fonds National de Recherche Scientifique [FRFS-WELBIO CR-2017S-03, FNRS CDR J.0068.19, FNRS-PDR T.0066.18]; Joint Programming Initiative on Antimicrobial Resistance [JPI-EC-AMR -R.8004.18]; Program ‘Actions de Recherche Concerté 2016–2021 and Fonds d’Encouragement à la Recherche (FER) of ULB; Fonds Jean Brachet, the Fondation Van Buuren and the ERC CoG DiStRes (Grant Agreement n° 864311) to A.G.P.; the European postdoctoral programme (Marie Skłodowska Curie COFUND action) to A.A..

## AUTHOR CONTRIBUTIONS

VH coordinated the study and drafted the manuscript with contributions from all authors. TK, AGP and VH designed experiments and analysed the data.TK, TB, SRAO, MR, KJT, OB, HT, AA, HTaman performed experiments. DR, RP, TT and AGP provided reagents. GCA performed bioinformatic analyses.

## SUPPLEMENTAL FIGURES AND TABLES

**Figure S1.**
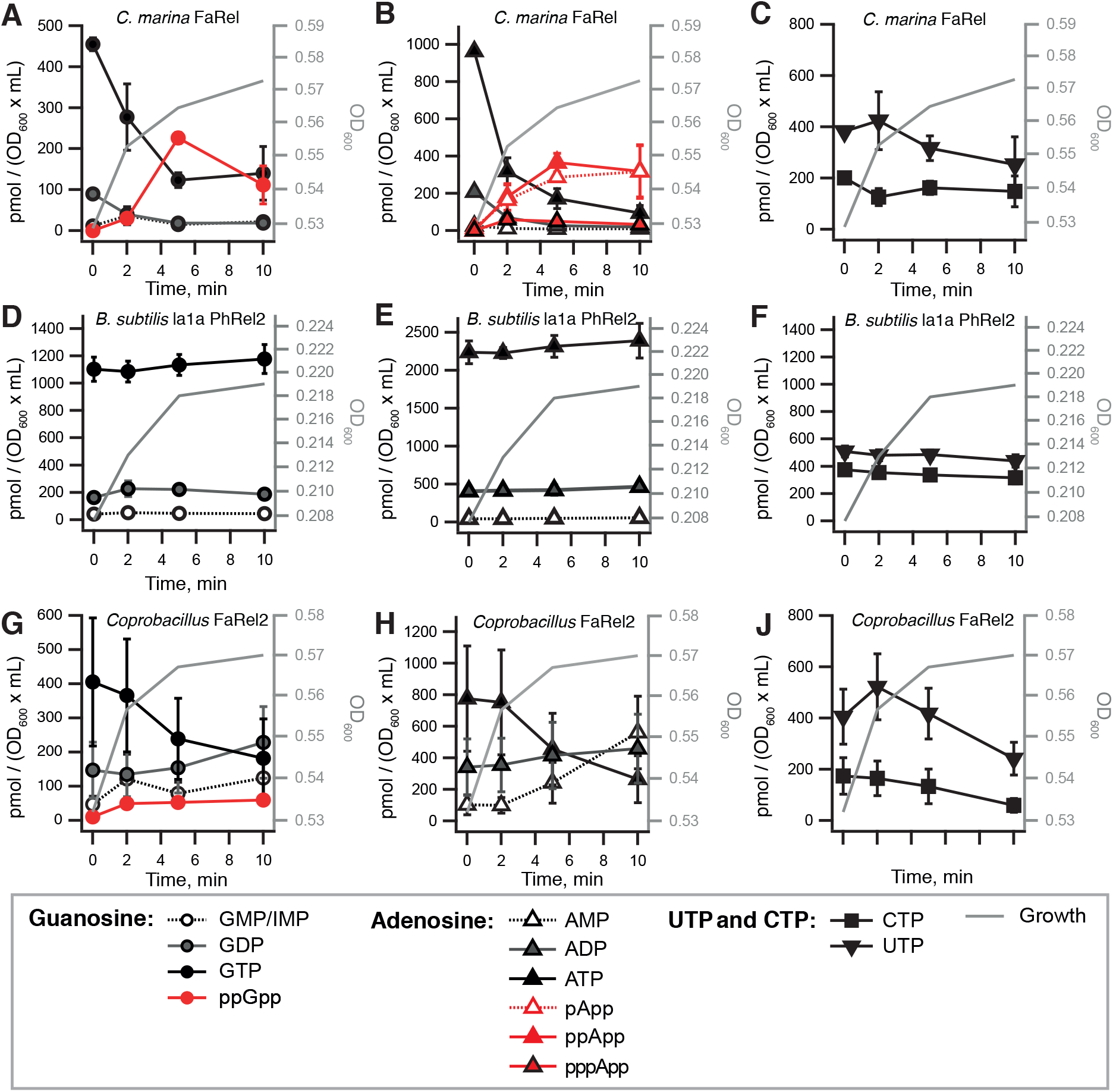
Nucleotide pools in *E. coli* BW25113 expressing *C. marina* FaRel, *B. subtilis* la1a PhRel2 and *Coprobacillus* sp. D7 FaRel2, related to Figure 1. Cell cultures were grown in defined minimal MOPS medium supplemented with 0.5% glycerol at 37 °C with vigorous aeration. The expression of toxic SAS RSHs was induced with 0.2% L-arabinose at the OD_600_ 0.5 (FaRel and FaRel2) or 0.2 (PhRel2). Intracellular nucleotides are expressed in pmol per OD_600_ • mL as per the insert. Error bars indicate the standard error of the arithmetic mean of biological replicates.

**Figure S2.**
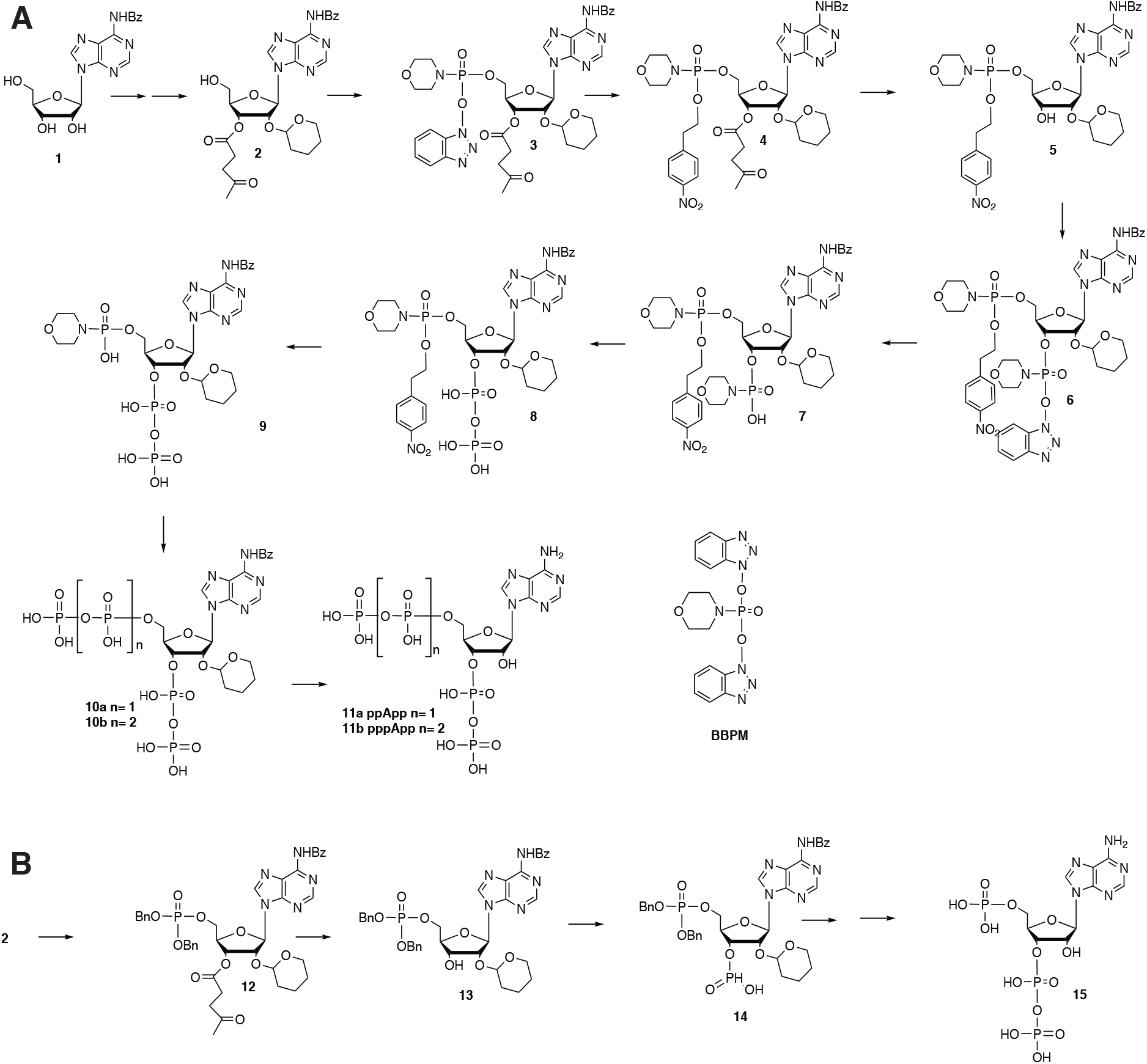
Synthesis of pppApp, ppApp and pApp, related to Figure 1. (**A**) Synthesis of ppApp and pppApp. N^6^-benzoyl adenosine (**1**) was used as a key starting material. Protected adenosine **2** reacted with reagent **BBPM** (bis(benzotriazolyl)phosphomorpholidate, see instert) affording intermediate **3**. Benzotriazolyl group of **3** was exchanged for 4-nitrophenylethyl group yielding **4**. Next, the 3’-levulinyl protecting group was removed (hydrazine, AcOH, pyridine). The resultant intermediate **5** was reacted again with BBPM affording **6**. Hydrolysis of benzotriazolyl group (Et_3_N, H_2_O, MeCN) provided morpholidate **7** that reacted with tributylamonium salt of phosphoric acid yielding 3’-diphosphate **8**. Removal of 4-nitrophenylethyl group (DBU, MeCN) afforded 5’-morpholidate **9** that upon reaction with tributylamonium salt of phosphoric or pyrophosphoric acid formed protected tetra-**10a** or pentaphosphate **10b**. The final products, ppApp (**11a**) and ppApp (**11b**), were obtained by removal of remaining protecting groups. Importantly, benzoyl group should be removed from nucleobase by treatment with aqueous ammonia first followed by removal of the THP group with 0.1N HCl (to avoid a nucleophilic attack of 2’-hydroxyl oxygen atom on phosphorus atom of 3’-phosphate moiety). (**B**) Synthesis of pApp. Dibenzyl phosphate was installed to 5’-position of protected adenosine **2** by reaction with dibenzyl diisopropylphosphoramidite under tetrazole catalysis followed by oxidation with 4-chloroperbenzoic acid affording **12**. Removal of levulinyl protecting group with hydrazine afforded intermediate **13** that was finally converted to pApp **14** using the same methodology as for the synthesis of ppApp.

**Figure S3.**
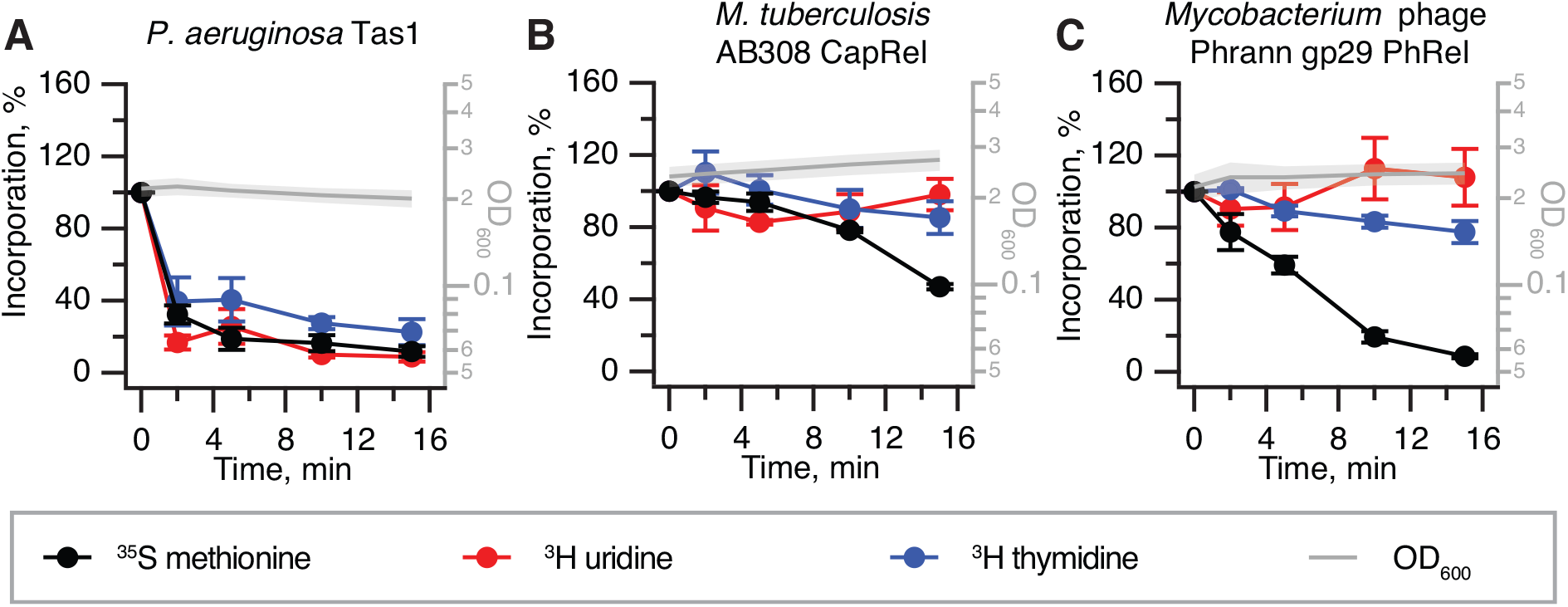
While expression of the (pp)pApp-synthesising *P. aeruginosa* Tas1 secretion system effector results in inhibition of transcription, translation and replication, the expression of *M. tuberculosis* AB308 CapRel or *Mycobacterium* phage Phrann PhRel toxSAS leads to specific inhibition of translation, related to Figure 1. Pulse-labelling assays following incorporation of ^35^S-methionine (black traces), ^3^H-uridine (red traces), and ^3^H-thymidine (blue traces). Expression of *P. aeruginosa* Tas1 (**B**), *M. tuberculosis* AB308 CapRel (**A**) or *Mycobacterium* phage Phrann PhRel (Gp29) (**c**) from the pBAD33-based constructs was induced by the addition of L-arabinose (final concentration 0.2%) to bacterial cultures in early exponential phase.

**Figure S4.**
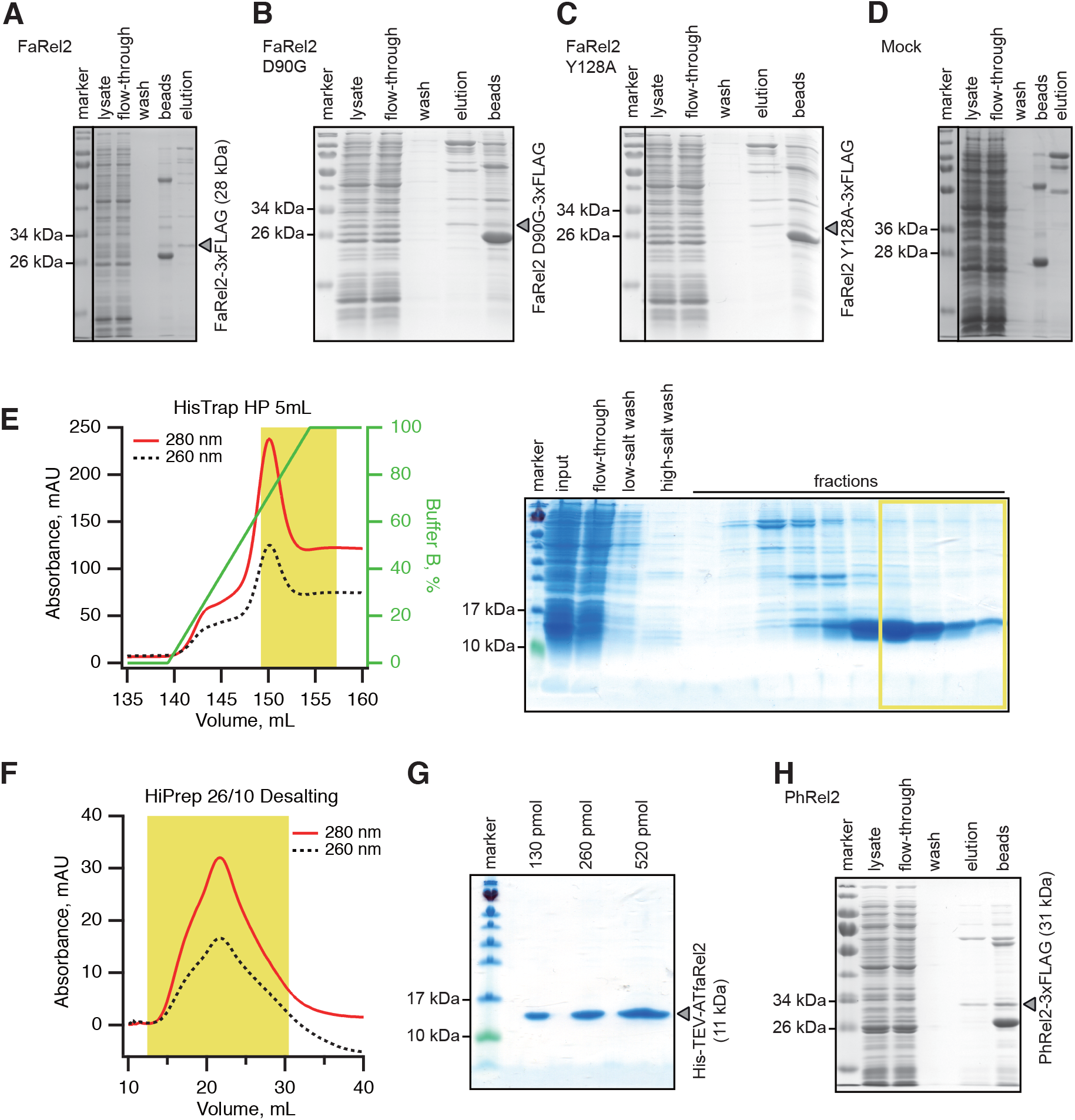
Protein purification of the FaRel2 toxSAS variants (A-D), ATfaRel2 antitoxin (E-G) and PhRel2 toxSAS, related to Figures 1, 2, 3 and S6. FLAG_3_-taged FaRel2 (**A**), FaRel2 D90A (**B**) and FaRel2 Y128A (**C**) proteins were immunoprecipitated with the anti-FLAG antibody and eluted with FLAG_3_ peptide. (**D**) Mock sample preparation from *E. coli* transformed with an empty vector (pBAD33) followed the same procedure as for toxin purification. Samples in each step were resolved by SDS-PAGE and visualised by Blue silver staining. (**E**) Cells expressing N-terminally His_6_-TEV-tagged ATfaRel2 were lysed and subjected to immobilised metal affinity chromatography (IMAC) using a HisTrap 5 mL HP column. The fraction corresponding to ATfaRel2 with the lowest contamination of other proteins (highlighted in yellow) was carried forward. Following the buffer exchange on HiPrep 10/26 desalting column (**F**), the fractions highlighted in yellow were pooled, concentrated, aliquoted, flash-frozen in liquid nitrogen and stored at –80 °C. (**G**) SDS-PAGE analysis of the purified ATfaRel2. (**H**) FLAG_3_-taged PhRel protein was immunoprecipitated with the anti-FLAG antibody and eluted with FLAG_3_ peptide.

**Figure S5.**
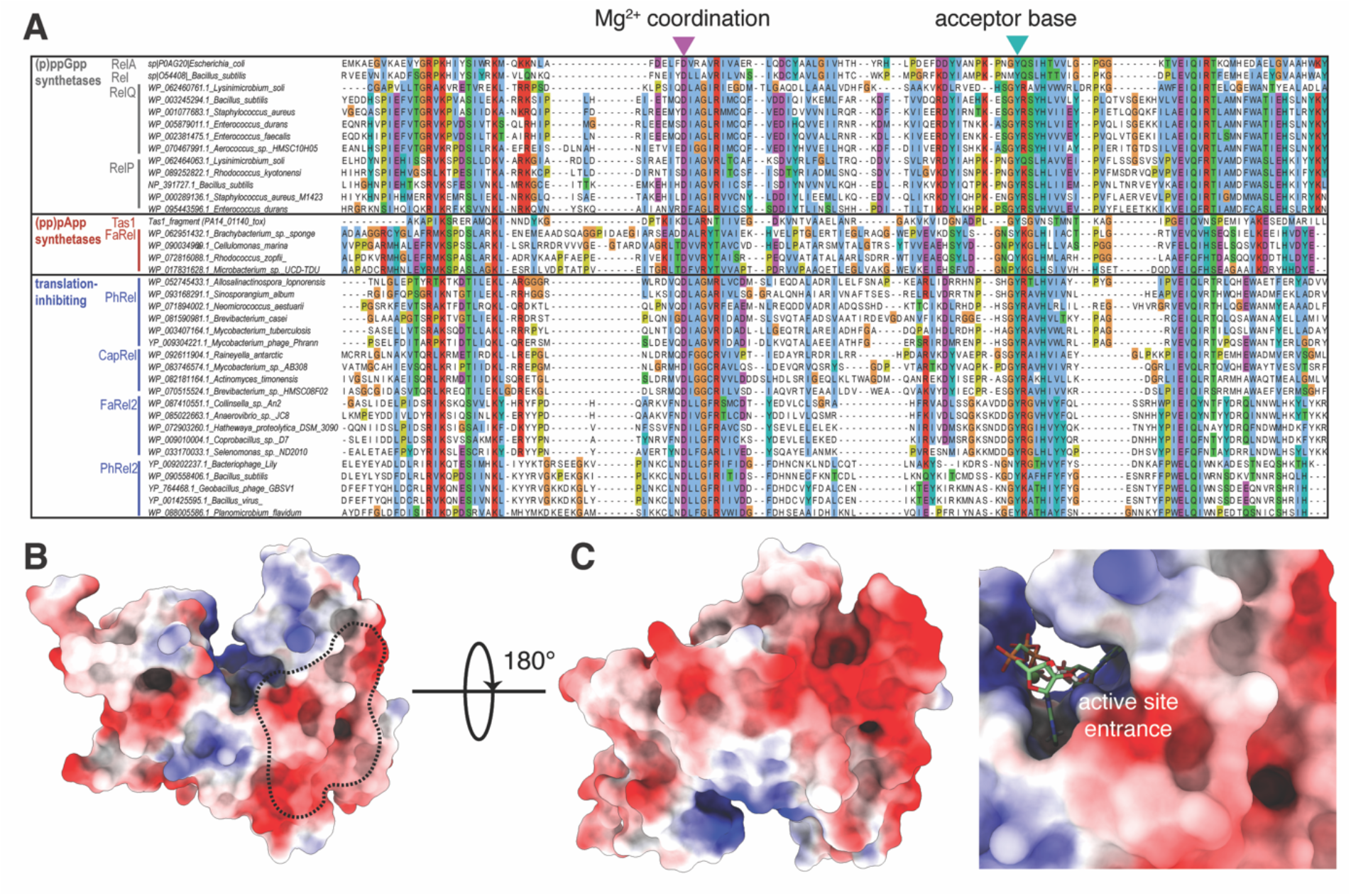
Sequence and structure of RSH SYNTH active sites, related to Figure 4. (**A**) Sequence alignment of the SYNTH domain of selected RSHs. Sequences are divided into groups based on their functional capabilities (left panel). Invariant functionally important positions that are substituted in *Coprobacillus* sp. D7 FaRel2 are indicated with triangles (D90A and Y128A shown in purple and turquoise respectively). (**B**) Surface charge distribution of *Staphylococcus aureus* RelQ (PDBID 6EWZ), the region equivalent to the tRNA-binding interface discovered in FaRel2 (highlighted in black) is highly negatively charged and likely precludes tRNA binding. (**C**) ATP and GDP binding sites are positively charged to stabilize the poly-phosphate groups of ATP and GDP/GTP.

**Figure S6.**
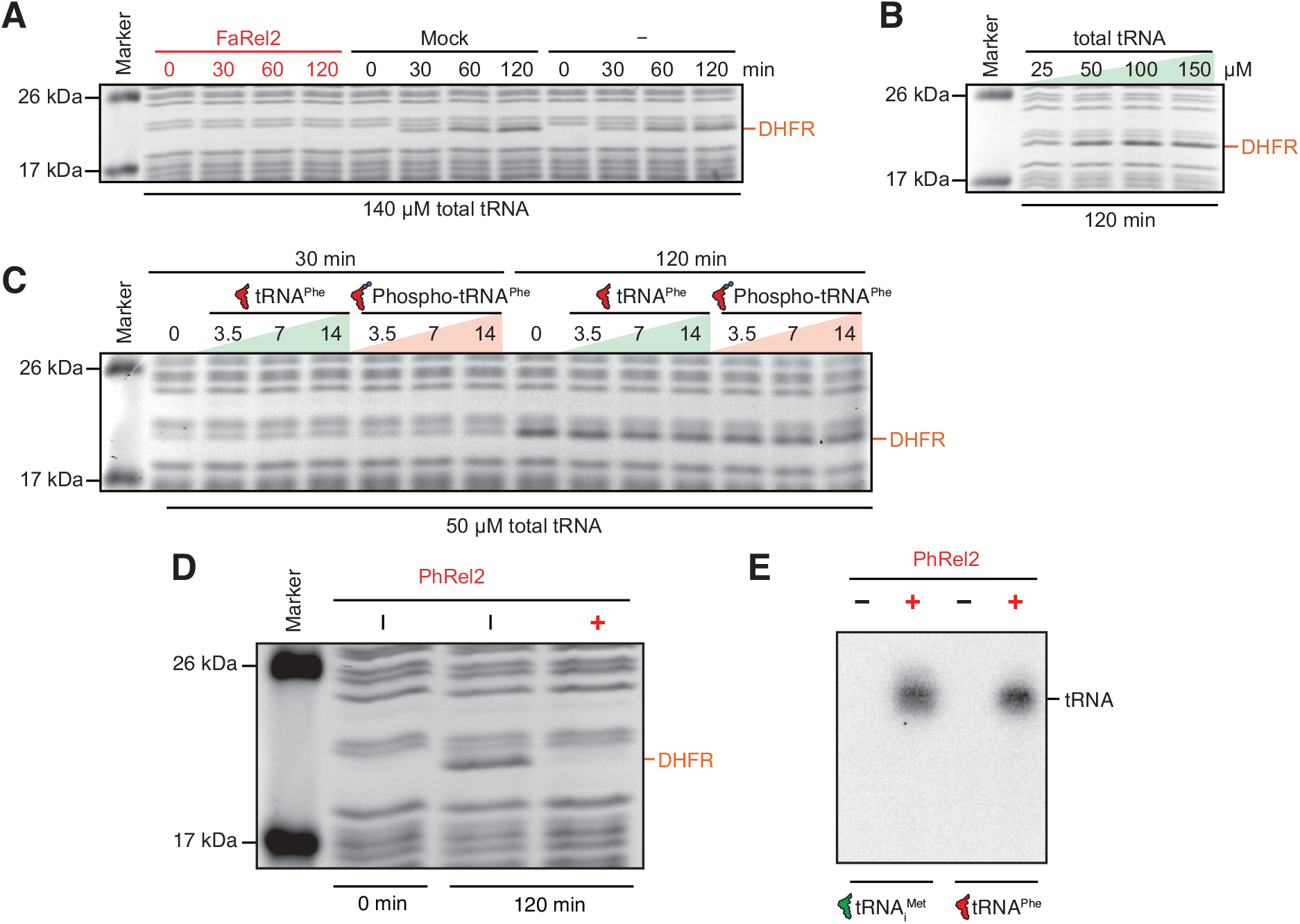
tRNA^Phe^ phosphorylated by FaRel2 does not inhibit translation in PURE system, related to Figures 1 and 2. (**A**) The cell-free protein synthesis reactions were supplemented with same volume of either 50 nM FaRel2-FLAG_3_ purified from *E. coli* FaRel2-FLAG_3_ expressing strain (FaRel2), mock protein preparation from *E. coli* strain transformed with an empty plasmid vector (mock) or HEPES:Polymix buffer pH 7.5 (–). (**B**) The concentration of total *E. coli* tRNA was titrated in PURE system and DHFR production was assessed by SDS PAGE at 120 minutes. (**C**) Phosphorylated elongator tRNA^Phe^ titrated into the PURE system containing 50 µM total tRNA. Cell-free synthesis reactions were performed at 37 °C, the duration of the reaction is indicated on the individual panels. (**D**) Addition of *B. subtilis* la1a PhRel2-FLAG_3_ abrogates production of DHFR in cell-free expression assays. (**E**) A reconstituted ^32^P transfer reaction using PhRel2and either tRNA_i_^fMet^ or tRNA^Phe^ as a substrate.

**Table S1.**
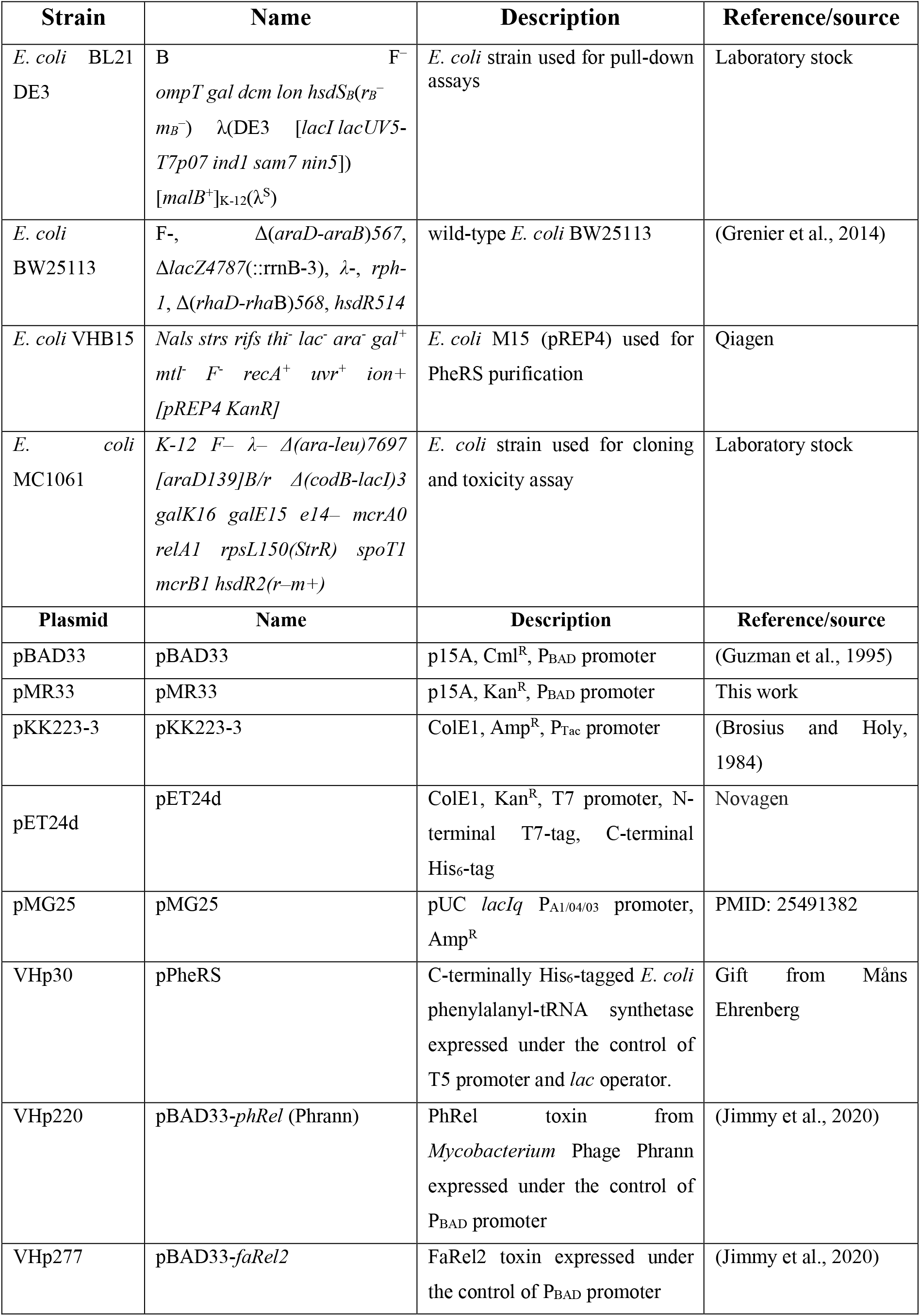
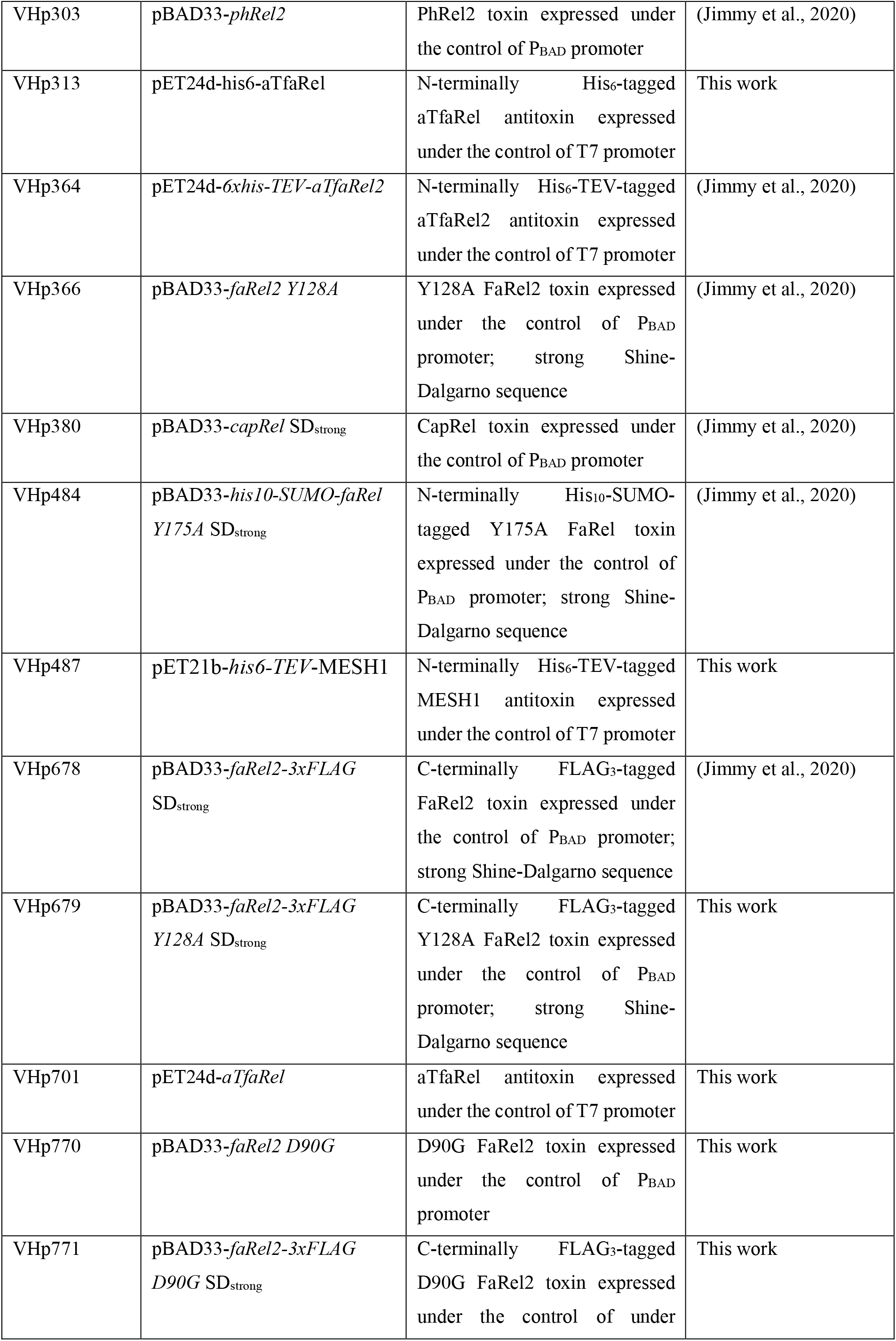
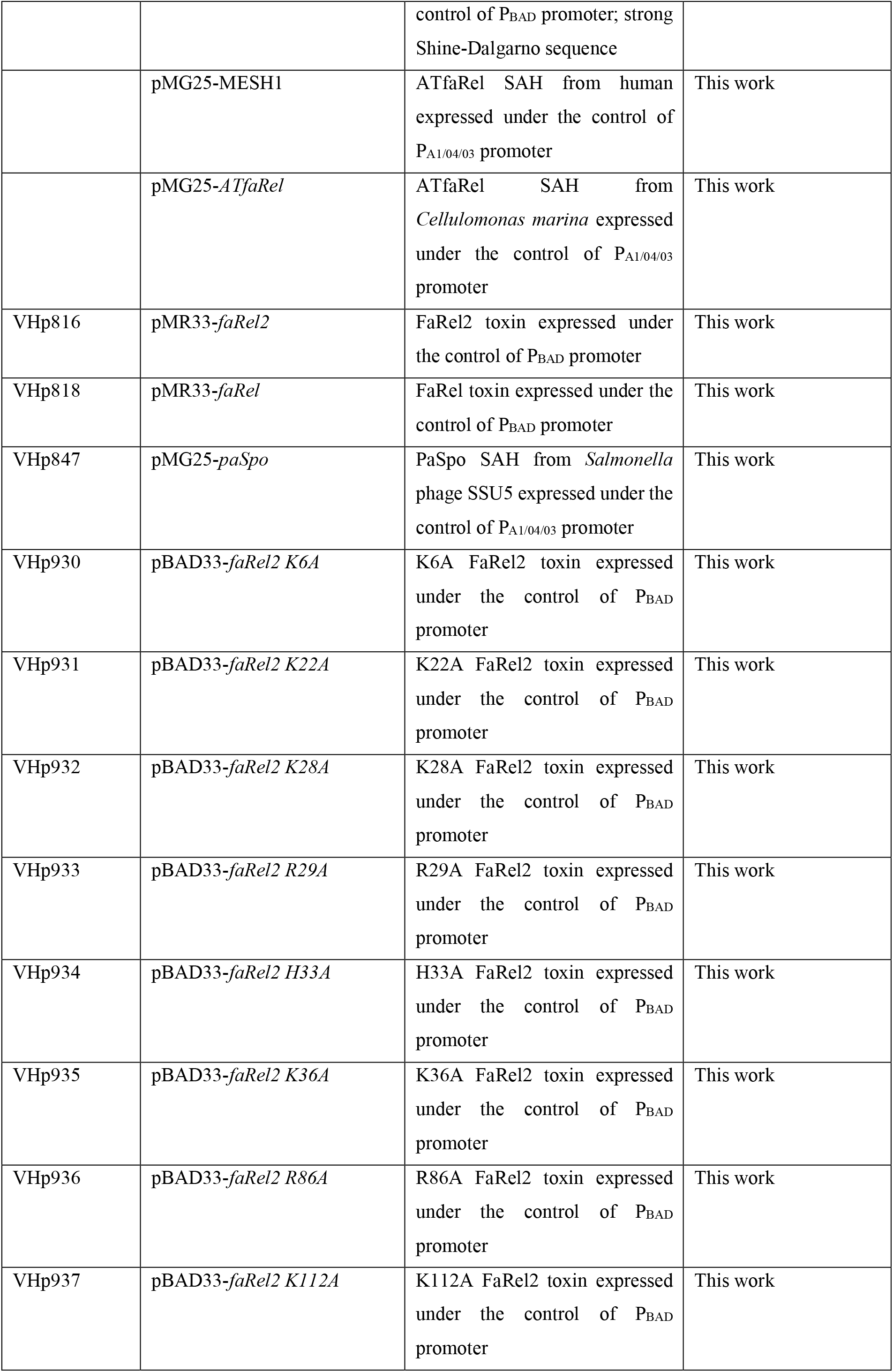
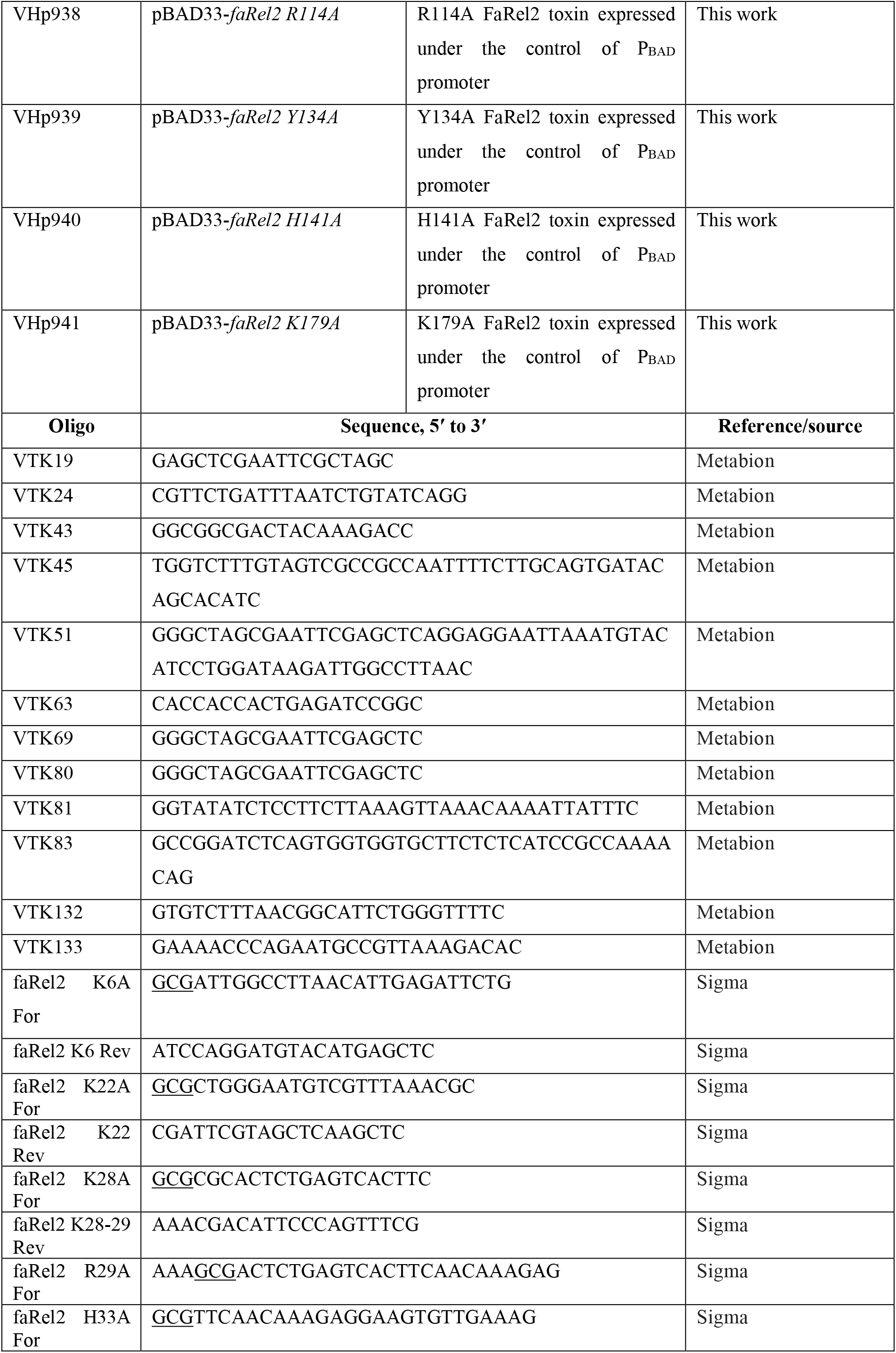
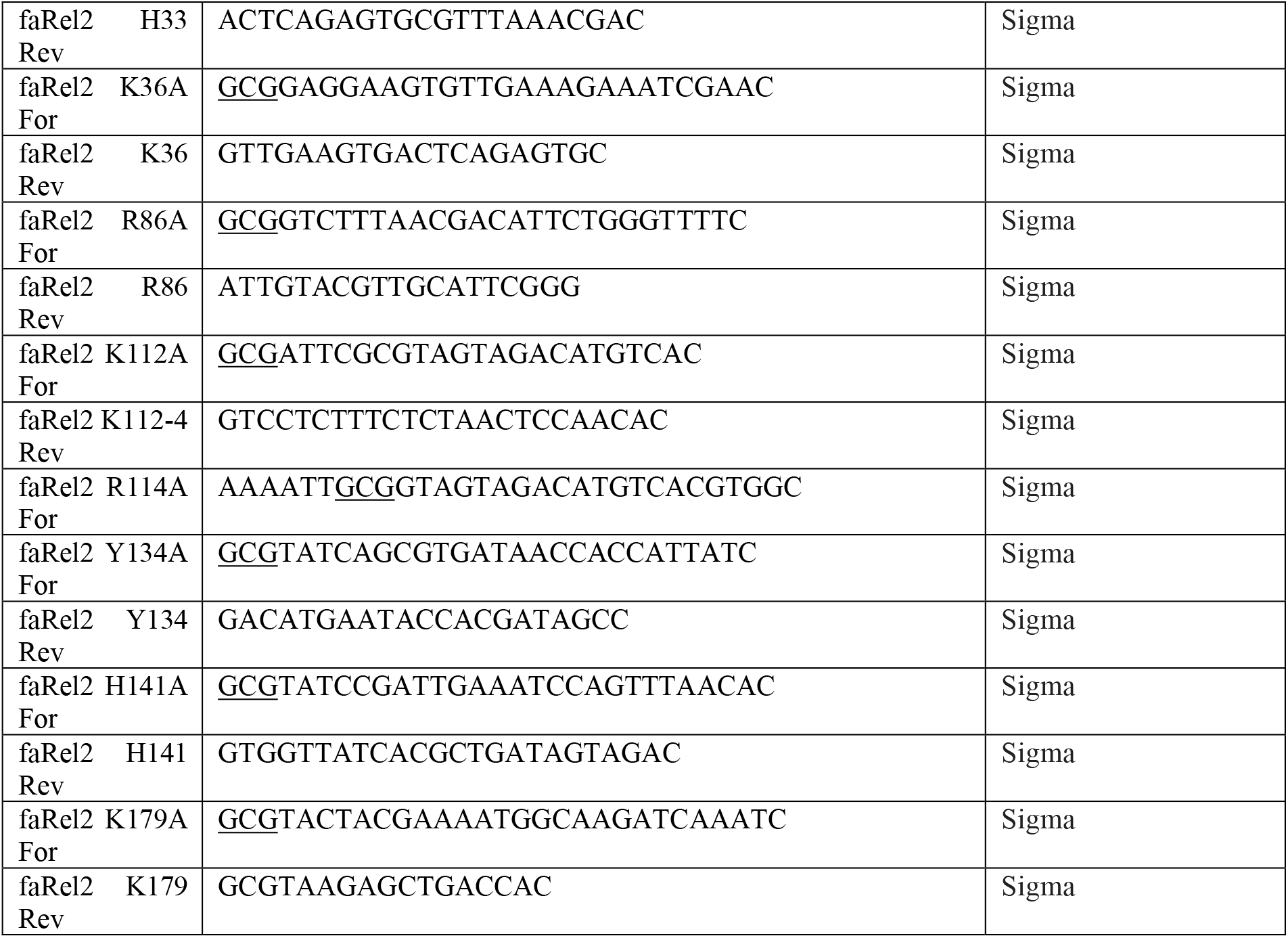
Strains, plasmids and oligonucleotide primers used in this study.

## METHODS

### Bacterial strains

Bacterial strains and plasmids as well as oligonucleotide primers used in the study are listed in **Table S1**.

### Multiple sequence alignment

Sequences were sequences extracted from the RSH database (Jimmy et al., 2020), aligned with MAFFT v7.164b with the L-ins-i strategy (Katoh and Standley, 2013), and alignments were visualised with Jalview (Waterhouse et al., 2009).

### Construction of plasmids

Oligonucleotides were synthesised by Metabion and Sigma. To construct the plasmids, DNA fragments were amplified by PCR and assembled by NEBuilder HiFi DNA Assembly Cloning Kit (NEB, E5520S).

To construct VHp770, DNA fragments were amplified by PCR using VHp277 as a template as well as sets of primers VTK69 and VTK133 or VTK19 and VTK132. To construct VHp701, VHp308(Jimmy et al., 2020) and pET24d were used as PCR templates with primer sets of VTK80 and VTK83 or VTK63 and VTK81 respectively. To generate VHp771, the DNA fragments were amplified by PCR using VHp678 as the template and sets of primers VTK69 and VTK133 or VTK19 and VTK132. To construct VHp679, VHp366 and VHp678 were used as PCR templates with primers VTK51 and VTK45 or VTK19 and VTK43, respectively. The identity of the constructed plasmids was confirmed through re-sequencing (LGC genomics).

VHp818 and VHp816 were constructed by sub-cloning *faRel* and *faRel2* from VHp307 (Jimmy et al., 2020) and VHp227(Jimmy et al., 2020), respectively, into pMR33 using the restriction enzymes SacI and HindIII. The constructed plasmids were validated through sequencing (LGC genomics). The point mutations to *faRel2* was done in the plasmid VHp277 background and introduced by amplifying the entire plasmid with divergent primers listed in **Table S1**. The forward primer introduced the desired amino-acid substitution in its unbound 5′ region. After PCR with Q5 polymerase (NEB), the product was treated with DpnI (NEB) to remove the template plasmid, purified trough a PCR purification column (Omega), phosphorylated with PNK (NEB) and ligated (NEB). The mixture was transformed into *E. coli* MC1061 and the resulting plasmids were verified by sequencing (Eurofins).

### Synthesis and characterisation of (pp)pApp

The synthesis of (p)ppApp (**Figure S2A**) followed the same procedure as for the preparation of pppGpp (Schattenkerk et al., 1985) but instead protected guanosine, N^6^-benzoyl adenosine (**1**) was used as the starting material. The final products ((p)ppApp, **11a** and **11b**) were purified by preparative reversed phase HPLC using linear gradient of methanol in 0.1M aqueous TEAB. Triethylammonium salt was converted to potassium salt by passing through small column with Dowex 50 in K^+^ phase, lyophilised from water and characterised by NMR and HR-MS. The synthesis of ppApp (**Figure S2B**) has been described in (Jimmy et al., 2020).

The synthesis of pApp followed a slightly different path. Dibenzyl phosphate was installed to the 5’-position of the protected adenosine **2** by reaction with dibenzyl diisopropylphosphoramidite under tetrazol catalysis, followed by oxidation with 4-chloroperbenzoic acid. Removal of the levulinyl protecting group was followed by installation of 3’-pyrophosphate using the same methodology as for the synthesis of ppApp.

#### ppApp 11a (I76DR_242P1)

^1^H NMR (500.2 MHz, D_2_O, ref(*t*BuOH) = 1.24 ppm): 4.21 – 4.27 (m, 2H, H-5); 4.59 (p, 1H, *J*_4’,3’_ = *J*_4’,5’_ = *J*_H,P_ = 2.9, H-4’); 4.88 (ddd, 1H, *J*_2’,1’_ = 6.5, *J*_2’,3’_ = 5.0, *J*_H,P_ = 1.3, H-2’); 4.98 (ddd, 1H, *J*_H,P_ = 8.3, *J*_3’,2’_ = 5.0, *J*_3’,4’_ = 2.9, H-3’); 6.22 (d, 1H, *J*_1’,2’_ = 6.5, H-1’); 8.28 (s, 1H, H-2);8.57 (s, 1H, H-8).

^13^C NMR (125.8 MHz, D_2_O, ref(*t*BuOH) = 32.43 ppm): 67.93 (d, *J*_C,P_ = 5.3, CH_2_-5’); 76.54 (d, *J*_C,P_ =4.5, CH-2’); 77.75 (d, *J*_C,P_ = 5.2, CH-3’); 86.41 (dd, *J*_C,P_ = 9.1, 3.8, CH-4’); 89.37 (CH-1’); 121.50 (C-5); 142.82 (CH-8); 152.17 (C-4); 155.73 (CH-2); 158.50 (C-6).

^31^P{^1^ H} NMR (202.5 MHz, D_2_O): -10.43 (d, *J* = 21.8, P_*α*_-3’); -10.33 (d, *J* = 20.7, P_*α*_-5’); -8.20 (bd, *J* = 20.7, P_*β*_-5’); -6.45 (bd, *J* = 21.8, P_*β*_-3’).

IR *v*_max_ (KBr) 3436 (vs, br), 3250 (m, br, sh), 3155 (m, br, sh), 1636 (m, br), 1578 (w, sh), 1475 (w, br, sh), 1337 (vw), 1301 (vw), 1220 (w, br), 1103 (w, br), 1074 (w, br), 972 (w, br), 921 (w, br), 797 (vw).

HR-MS(ESI^-^) For C_10_H_16_O1_6_N_5_P_4_ (M-H)^-^ calcd 585.95480, found 585.95518.

#### pppApp 11b (I76DR_265P1)

^1^H NMR (500.2 MHz, D_2_O, ref(*t*BuOH) = 1.24 ppm): 4.25 (ddd, 1H, *J*_gem_ = 11.7, *J*_H,P_ = 4.9, *J*_5’b,4’_ = 2.8, H-5’a); 4.28 (ddd, 1H, *J*_gem_ = 11.7, *J*_H,P_ = 5.7, *J*_5’a,4’_ = 2.8, H-5’a); 4.65 (p, 1H, *J*_4’,3’_ = *J*_4’,5’_ = *J*_H,P_ = 2.8, H-4’); 4.87 (ddd, 1H, *J*_2’,1’_ = 7.0, *J*_2’,3’_ = 5.2, *J*_H,P_ = 1.3, H-2’); 4.96 (bm, 1H, H-3’); 6.19 (d, 1H, *J*_1’,2’_ = 7.0, H-1’); 8.27 (s, 1H, H-2); 8.56 (s, 1H, H-8).

^13^C NMR (125.8 MHz, D_2_O, ref(*t*BuOH) = 32.43 ppm): 68.38 (d, *J*_C,P_ = 5.2, CH_2_-5’); 76.66 (d, *J*_C,P_ = 4.9, CH-2’); 78.21 (d, *J*_C,P_ = 6.0, CH-3’); 86.45 (dd, *J*_C,P_ = 8.7, 2.9, CH-4’); 89.08 (CH-1’); 121.48 (C-5); 142.81 (CH-8); 152.25 (C-4); 155.66 (CH-2); 158.45 (C-6).

^31^P{^1^H} NMR (202.5 MHz, D_2_O): -22.14 (t, *J* = 19.1, P_*β*_-5’); -10.83 (d, *J* = 20.6, P_*α*_-3’); -10.63 (d, *J* = 19.1, P_*α*_-5’); -9.65 (bd, *J* = 19.1, P_*γ*_-5’); -8.63 (bd, *J* = 20.6, P_*β*_-3’).

HR-MS(ESI^-^) For C_10_H_17_O_19_N_5_P_5_ (M-H)^-^ calcd 665.92113, found 665.91960.

#### pApp (I76DR_215P1)

^1^H NMR (500.2 MHz, D_2_O, ref(*t*BuOH) = 1.24 ppm): 4.12 (ddd, 1H, *J*_gem_ = 11.8, *J*_H,P_ = 4.6, *J*_5’b,4’_ = 2.8, H-5’a); 4.16 (ddd, 1H, *J*_gem_ = 11.8, *J*_H,P_ = 4.8, *J*_5’a,4’_ = 2.8, H-5’a); 4.59 (dt, 1H, *J*_4’,3’_ = 3.2, *J*_4’,5’_ = 2.8, H-4’); 4.85 (ddd, 1H, *J*_2’,1’_ = 6.2, *J*_2’,3’_ = 5.1, *J*_H,P_ = 1.3, H-2’); 4.96 (ddd, 1H,

*J*_H,P_ = 8.5, *J*_3’,2’_ = 5.1, *J*_3’,4’_ = 3.2, H-3’); 6.19 (d, 1H, *J*_1’,2’_ = 6.2, H-1’); 8.26 (s, 1H, H-2); 8.53 (s, 1H, H-8).

^13^C NMR (125.8 MHz, D_2_O, ref(*t*BuOH) = 32.43 ppm): 67.01 (d, *J*_C,P_ = 4.7, CH_2_-5’); 76.58 (d, *J*_C,P_ =4.6, CH-2’); 77.57 (d, *J*_C,P_ = 5.4, CH-3’); 86.34 (dd, *J*_C,P_ = 8.9, 4.0, CH-4’); 89.55 (CH-1’); 121.53 (C-5); 142.82 (CH-8); 152.09 (C-4); 155.71 (CH-2); 158.49 (C-6).

^31^P{^1^H} NMR (202.5 MHz, D_2_O): -10.69 (d, *J* = 21.1, P_*α*_-3’); -7.94 (d, *J* = 21.2, P_*β*_-3’); 1.51 (s, P-5’).

HR-MS(ESI^-^) For C_10_H_15_O_13_N_5_P_3_ (M-H)^-^ calcd 505.98847, found 505.98861.

### HPLC-based nucleotide quantification

*E. coli* strain BW25113 (Grenier et al., 2014) was transformed with RSH-expressing plasmids (pMR33-faRel, pMR33-faRel2 or pBAD33-phRel2) as well as empty pKK223-3 vector. The starter cultures were pre-grown overnight at 37 °C with vigorous shaking (200 rpm) in Neidhardt MOPS minimal media (Neidhardt et al., 1974) supplemented with 1 µg/mL thiamine, 1% glucose, 100 µg/mL carbenicillin as well as either 20 µg/mL chloramphenicol (pBAD33--phRel2) or 25 µg/mL kanamycin (pMR33-faRel and pMR33-faRel2). The overnight cultures were diluted to OD_600_ 0.05 in 115 mL of pre-warmed medium MOPS supplemented with 0.5% glycerol as carbon source and grown until OD_600_ ≈ 0.5 (pMR33-faRel and pMR33-faRel2) or OD_600_ ≈ 0.2 (pBAD33-phRel2) at 37 °C, 200 rpm. At this point 0.2% arabinose was added to induce the expression of the toxin. 26 mL samples were collected for HPLC analyses at 0, 2, 5 and 10 minutes after the addition of arabinose and IPTG. Nucleotide extraction and HPLC analyses were performed as described previously (Varik et al., 2017). The OD_600_ measurements were performed in parallel with collection of the samples for HPLC analyses.

## Metabolic labelling with ^35^S-methionine, ^3^H-uridine or ^3^H-thymidine

Overnight cultures in defined Neidhardt MOPS minimal media (Neidhardt et al., 1974) supplemented with 1% glucose, 0.1% casamino acids and as well as appropriate antibiotics were inoculated with single colonies of *E. coli* BW25113 cells freshly transformed with pBAD33-based plasmid for L-arabinose-inducible RSH expression as well as the empty pKK223-3 vector. After overnight incubation at 37 °C with shaking at 180 rpm, the cultures were diluted to an OD_600_ of 0.05 in 15 mL MOPS minimal media supplemented with 19 amino acids (25 µg/mL, final concentration) but lacking methionine, 0.5% glycerol, as well as appropriate antibiotics. The cultures were grown at 37 °C until an OD_600_ of 0.2-0.3 in a water bath with shaking (200 rpm), and expression of toxins was induced with 0.2% L-arabinose. For a zero-point 1 mL of culture was taken and mixed with either 4.35 µCi ^35^S-methionine (Perkin Elmer), 0.65 µCi ^3^H-uridine (Perkin Elmer) or 2 µCi ^3^H-thymidine (Perkin Elmer) immediately before induction. Simultaneously, another 1 mL of culture was taken for OD_600_ measurements. Samples were collected at 2, 5, 10 and 15 minutes post-induction and processed as described above. The incorporation of radioisotopes was quenched 8 minutes after addition of the isotope, by the addition of 200 µL ice-cold 50% trichloroacetic acid (TCA). Samples were filtered through GF/C filters (Whatman) prewashed with 5% TCA, followed by washing twice with 5 mL of ice-cold TCA and, finally, twice with 5 mL of 95% EtOH. The filters were dried at least for two hours at room temperature and the radioactivity was quantified by scintillation using EcoLite Liquid Scintillation Cocktail scintillation cocktail (5 mL per vial, MP Biomedicals, 15 minutes shaking with filters prior to counting) using a TRI-CARB 4910TR 100 V scintillation counter (PerkinElmer).

### Toxicity validation assays

The experiments were performed as described earlier (Jimmy et al., 2020). The assays were performed on LB medium (Lennox) plates (VWR). We used *E. coli* BW25113 strain co-transformed with two different plasmid systems for controllable expression of toxins and antitoxin.

First, we used a combination of pKK223-3 (medium copy number, ColE1 origin of replication, Amp^R^, antitoxins expressed under the control of P_Tac_ promoter (Brosius and Holy, 1984)) and pBAD33 harbouring toxin genes (medium copy number, p15A origin of replication, Cml^R^, toxins expressed under the control of P_BAD_ promoter (Guzman et al., 1995)) (**Figure 1D** and **Figure 4B**). The cells were grown in liquid LB medium (BD) supplemented with 100 µg/mL carbenicillin (AppliChem) and 20 µg/mL chloramphenicol (AppliChem), 30 mM K_2_HPO_4_/KH_2_PO_4_ (pH 7.4) as well as 1% glucose (repression conditions). Serial ten-fold dilutions were spotted (5 µl per spot) on solid LB plates containing carbenicillin and chloramphenicol in addition to either 1% glucose (repressive conditions), or 0.2% arabinose combined with 1 mM IPTG (induction conditions). Plates were scored after an overnight incubation at 37 °C. Sequences were codon-optimised for expression in *E. coli*.

Second, we used pMG25 (high copy number, ColE1 origin of replication (pUC), Amp^R^, antitoxin expressed under the control of IPTG inducible P_A1/04/03_ promoter (Jaskolska and Gerdes, 2015)) and pBAD-based pMR33 (this work) harbouring toxin genes (medium copy number, p15A origin of replication, Kan^R^, toxins expressed under the control of P_BAD_ promoter) (**Figure 3A**). The cells were grown in liquid LB medium (BD) supplemented with 0.2% glucose (repression conditions), 100 µg/mL carbenicillin (AppliChem) and 50 µg/mL kanamycin (AppliChem). Serial dilutions and spotting were performed as described above using solid LB plates supplemented with 0.2% arabinose as well as 100 µg/mL carbenicillin (AppliChem) and 50 µg/mL kanamycin (AppliChem).

### Protein expression and purification

The *Coprobacillus* sp. D7 C-terminally FLAG_3_-tagged FaRel2 (FaRel2-FLAG_3_) was overexpressed in freshly transformed *E. coli* BL21 DE3 co-transformed with the VHp701 plasmid encoding the non-tagged SAH aTfaRel antitoxin under the pET promoter. Fresh transformants were inoculated to a final OD_600_ of 0.04 in the LB medium (800 mL) supplemented with 100 µg/mL kanamycin and 20 µg/mL chloramphenicol. The cultures were grown at 37 °C until an OD_600_ of 0.3, the antitoxin was pre-induced with 0.1 mM IPTG (final concentration) for one hour and the toxin was induced with 0.2% arabinose (final concentration) for an additional one hour at 37 °C. The cells were collected by centrifugation (8,000 rpm, 10 minutes at 4 °C, JLA-10.500 Beckman Coulter rotor), dissolved in 4 mL of cell suspension buffer (20 mM HEPES:KOH pH 7.5, 95 mM KCl, 5 mM NH_4_Cl, 0.5 mM CaCl_2_, 8 mM putrescine, 1 mM spermidine, 5 mM Mg(OAc)_2_, 1 mM DTT and cOmplete protease inhibitor (Mini, EDTA-free from Roche)). The cell suspension was divided to 1 mL aliquots, and 200 µl of pre-chilled zirconium beads (0.1 mm) were added in the aliquots. Cellular lysates were prepared by a FastPrep homogeniser (MP Biomedicals) (four 20 seconds pulses at speed 4.5 mp per second with chilling on ice for 2 minutes between the cycles) and clarified by centrifugation at 21,000 g for 20 minutes at 4 °C. The supernatant was carefully collected, avoiding the lipid layer and cellular pellet.

30 mg of total protein (as determined by Bradford assay) of each sample was mixed with 100 µL of ANTI-FLAG M2 Affinity Gel (Sigma-Aldrich) and mixed by rotation for 2 hours at 4 °C. The mixture was loaded on a Micro Bio-Spin Chromatography Column (Bio-Rad) and flow-through was collected. The gel in the column was washed five times with 1 mL of cell suspension buffer supplemented with 10% glycerol, and the fraction at final wash was collected. The gel was mixed with 300 µL of cell suspension buffer supplemented with 10% glycerol as well as 0.1 mg/mL Poly FLAG Peptide lyophilised powder (Biotool) in the column by rotation for 40 min at 4 °C. The elution fraction was passed through the column by spinning down, and was collected in Eppendorf tube. After this elution step, the gel was suspended with 1x sample buffer (50 mM Tris:HCl pH 6.8, 2% SDS, 0.01% bromophenol blue, 10% glycerol, 10 mM DTT and 2% beta-mercaptoethanol) and collected. 0.5 µL of cell lysate, 0.5 µL of flowthrough, 8 µL of wash, 8 µL of elution fractions and 10 µL of gel suspension were resolved on 15% SDS-PAGE gel.

The SDS-PAGE gel was fixed with fixing solution (50% ethanol and 2% phosphoric acid) for 5 min at room temperature, washed with water for 20 minutes at room temperature twice, and stained with “blue silver” solution(Candiano et al., 2004) (0.12% Brilliant Blue G250 (Sigma-Aldrich, 27815), 10% ammonium sulfate, 10% phosphoric acid, and 20% methanol) overnight at room temperature. After washing with water for 3 hours at room temperature, the gel was analysed on an ImageQuant LAS 4000 (GE Healthcare) imaging system (**Figure S4A,D**). The concentration of FaRel2-FLAG_3_ was quantified on SDS-PAGE gels by ImageJ(Schneider et al., 2012) using pure ATfaRel2 as a standard.

For Western blotting, the proteins resolved by similar electrophoresis were transferred to 0.2 µm nitrocellulose membrane (BioTrace™ NT, Pall) using Trans-Blot® TurboTM Transfer System (Bio-Rad). To detect FLAG_3_-tagged protein, the membrane was blocked in PBS-T (1xPBS supplemented with 0.05% Tween-20) with 5% w/v nonfat dry milk at room temperature for one hour, and first antibody incubation was performed for overnight at 4 °C in PBS-T anti-Flag M2 (Sigma-Aldrich F1804, batch number #SLCD3524; 1:5000 dilution). After three 5-minute washes in fresh PBS-T, second antibody incubations were performed for one hour at room temperature in PBS-T with Goat anti-Mouse IgG-HRP (Agrisera AS11 1772, batch number #810-103-040; 1:4,000 dilution). Tagged-proteins were detected on an ImageQuant LAS 4000 (GE Healthcare) imaging system using WesternBright Quantum HRP substrate (Advansta).

The C-terminally FLAG_3_-tagged *B. subtilis* la1a PhRel2 (PhRel2-FLAG_3_) was overexpressed in freshly transformed *E. coli* BW25113 co-transformed with the VHp847 plasmid encoding the PaSpo SAH antitoxin under the IPTG-inducible promoter (P_A1/04/03_ promoter). Fresh transformants were inoculated to a final OD_600_ of 0.05 in the LB medium (800 mL) supplemented with 100 µg/mL carbenicillin, 20 µg/mL chloramphenicol and 50 µM IPTG. The cultures were grown at 37 °C until an OD_600_ of 0.5, the toxin was induced with 0.2% arabinose (final concentration) for an additional 3 hours at 37 °C. The cells were collected by centrifugation (8,000 rpm, 10 minutes at 4 °C, JLA-10.500 Beckman Coulter rotor), dissolved in 4 mL of cell suspension buffer (20 mM HEPES:KOH pH 7.5, 95 mM KCl, 5 mM NH_4_Cl, 0.5 mM CaCl_2_, 8 mM putrescine, 1 mM spermidine, 5 mM Mg(OAc)_2_, 1 mM DTT, and cOmplete, Mini, EDTA-free (Roche)). The cell suspension was divided to 1 mL aliquots, and 200 µl of pre-chilled zirconium beads (0.1 mm) were added in the aliquots. Cellular lysates were prepared by a FastPrep homogeniser (MP Biomedicals) (four 20 seconds pulses at speed 4.5 mp per second with chilling on ice for 2 minutes between the cycles) and clarified by centrifugation at 21,000 g for 20 minutes at 4 °C. The supernatant was carefully collected, avoiding the lipid layer and cellular pellet.

30 mg of total protein (as determined by Bradford assay) of each sample was mixed with 100 µL of ANTI-FLAG M2 Affinity Gel (Sigma-Aldrich) and mixed by rotation for 2 hours at 4 °C. The mixture was loaded on a Micro Bio-Spin Chromatography Column (Bio-Rad) and flow-through was collected. The gel in the column was washed with 1 mL of cell suspension buffer including 1 M KCl five times and 1 mL of cell suspension buffer supplemented with 10% glycerol five times, and the fraction at final wash was collected. The gel was mixed with 300 µL of cell suspension buffer supplemented with 10% glycerol and 0.1 mg/mL Poly FLAG Peptide lyophilised powder (Biotool) in the column by rotation for 40 min at 4 °C. The elution fraction was passed through the column by spinning down, and was collected in Eppendorf tube. After this elution step, the gel was suspended with 1x sample buffer (50 mM Tris:HCl pH 6.8, 2% SDS, 0.01% bromophenol blue, 10% glycerol, 10 mM DTT and 2% beta-mercaptoethanol) and collected. 0.5 µL of cell lysate, 0.5 µL of flowthrough, 8 µL of wash, 8 µL of elution fractions and 10 µL of gel suspension were resolved on 10% SDS-PAGE gel.

*Coprobacillus* sp. D7 N-terminally His_6_-TEV-tagged ATfaRel2 was overexpressed in freshly transformed *E. coli* BL21(DE3) with VHp364. Fresh transformants were inoculated to a final OD_600_ of 0.05 in the LB medium (800 mL) supplemented with 100 µg/mL kanamycin. The cultures were grown at 37 °C until an OD_600_ of 0.5, induced with 0.4 mM IPTG (final concentration) and grown for an additional one hour at 30 °C. The cells were harvested by centrifugation and resuspended in buffer A (300 mM NaCl, 10 mM imidazole, 10% glycerol, 4 mM β-mercaptoethanol, 25 mM HEPES:KOH pH 8.0) supplemented with 0.1 mM PMSF and 1 U/mL of DNase I, and lysed by one passage through a high-pressure cell disrupter (Stansted Fluid Power, 150 MPa). Cell debris was removed by centrifugation (25,000 rpm for 1 hour) and clarified lysate was taken for protein purification. Clarified cell lysate was filtered through a 0.22 µm syringe filter and loaded onto a HisTrap 5 mL HP column pre-equilibrated in buffer A. The column was washed with 5 column volumes (CV) of buffer A and following buffer B (1 M NaCl, 10 mM imidazole, 10% glycerol, 4 mM β-mercaptoethanol, 25 mM HEPES:KOH pH 8.0), and the protein was eluted using a linear gradient (3 CV with 0-100%) of buffer C (300 mM NaCl, 300 mM imidazole, 10% glycerol, 4mM β-mercaptoethanol, 25 mM HEPES:KOH pH 8.0). Fractions enriched in ATfaRel2 (≈60% buffer C) were pooled totaling approximately 8 mL (**Figure S4E**). The sample was applied on a HiPrep 10/26 desalting column (GE Healthcare) pre-equilibrated with storage buffer (buffer D; 300 mM KCl, 10% glycerol, 4 mM β-mercaptoethanol, 25 mM HEPES:KOH pH 8.0). The fractions containing ATfaRel2 were collected (about 8 mL in total, (**Figure S4F**) and concentrated on an Amicon Ultra (Millipore) centrifugal filter device with a 3 kDa cut-off. The purity of protein preparations was assessed by SDS-PAGE (**Figure S4G**). Protein preparations were aliquoted, frozen in liquid nitrogen and stored at –80 °C. Individual single-use aliquots were discarded after the experiment.

For purification of RelA *E. coli* BL21 DE3 harbouring pET24d:his10-SUMO-relA expression constructs were grown, induced, harvested and lysed as described earlier (Kudrin et al., 2018). Protein purification was performed as previously described (Turnbull et al., 2019).

C-terminally His_6_-tagged *E. coli* phenylalanyl-tRNA synthetase (PheRS) was overexpressed in *E. coli* BL21 (DE3). Fresh transformants were used to inoculate 3 L cultures of LB medium supplemented with 100 µg/mL ampicillin. The cultures were grown at 37 °C until an OD_600_ of 0.5, protein expression was induced with 1 mM IPTG (final concentration) and then the cultures were grown overnight at 37 °C. The cells were harvested by centrifugation, resuspended in buffer E (150 mM NaCl, 5 mM MgCl_2_, 20 mM imidazole, 1 mM β-mercaptoethanol, 20 mM

Tris:HCl pH 7.5) supplemented with 1 mM PMSF, and lysed by one passage through a high-pressure cell disrupter. Cell debris was removed by centrifugation and clarified lysate was taken for protein purification. Clarified cell lysate was filtered through a 0.22 µm syringe filter and loaded onto a HisTrap 5 mL HP column pre-equilibrated in buffer E. The column was washed with 20 column volumes (CV) of buffer F (1 M NaCl, 5 mM MgCl_2_, 20 mM imidazole, 1 mM β-mercaptoethanol, 20 mM Tris:HCl pH 7.5) and following buffer E (5 CV), and the protein was eluted using a linear gradient (30 CV with 0-100%) of buffer G (150 mM NaCl, 5 mM MgCl_2_, 500 mM imidazole, 1 mM β-mercaptoethanol, 20 mM Tris:HCl pH 7.5). Fractions enriched in PheRS (≈45% buffer G) were pooled. The sample was applied on a HiPrep 10/26 desalting column (GE Healthcare) pre-equilibrated with storage buffer (buffer H; 100 mM KCl, 2 mM MgCl_2_, 10% glycerol, 6 mM β-mercaptoethanol, 20 mM Tris:HCl pH 7.5). The fractions containing PheRS were collected. The protein was aliquoted, aliquots plunge-frozen in liquid nitrogen and stored at –80 °C. Individual single-use aliquots were discarded after the experiment.

Human MESH1 (VHp487, pET21b-His6-TEV-MESH1) was overexpressed in freshly transformed *E. coli* BL21 DE3 Rosetta (Novagen). Transformants were inoculated to a final OD_600_ of 0.05 in LB medium (400 mL×2) supplemented with 100 µg/mL Carbenicillin. The cultures were grown at 37 °C until an OD_600_ of 0.5, induced with 1 mM IPTG (final concentration) and grown for an additional 2 hours at 30 °C. The cells were harvested by centrifugation and resuspended in 20 ml of resuspension buffer (buffer I; 1 M KCl, 5 mM MgCl_2_, 1 mM β-mercaptoethanol, 50 mM Tris:HCl pH 8.0) supplemented with 0.1 mM PMSF, 1 mg/ml lysozyme and 1 U/mL of DNase I and incubated on ice for 30 min. After adding 10 mL of lysis buffer (buffer J; 500 mM KCl, 500 mM NaCl, 1% glycerol, 1 mM β-mercaptoethanol, 50 mM Tris:HCl pH 8.0), cells were lysed by one passage through a high-pressure cell disrupter (Stansted Fluid Power, 150 MPa), cell debris was removed by centrifugation (25,000 rpm for 40 min, JA-25.50 Beckman Coulter rotor) and clarified lysate was taken for protein purification.

Clarified cell lysate was filtered through a 0.2 µm syringe filter and loaded onto the HisTrap 1 mL HP column pre-equilibrated in buffer K (500 mM KCl, 500 mM NaCl, 10 mM MgCl_2_, 1 mM β-mercaptoethanol, 0.002% mellitic acid, 15 mM imidazole, 50 mM Tris:HCl pH 8.0). The column was washed with 5 CV of buffer K, and the protein was eluted with a linear gradient (20 CV, 0-100% buffer L) of buffer L (500 mM KCl, 500 mM NaCl, 10 mM MgCl_2_, 1 mM β-mercaptoethanol, 0.002% mellitic acid, 500 mM imidazole, 50 mM Tris:HCl pH 8.0). Fractions most enriched in His_6_-TEV-MESH1 (≈50-60% buffer B) were pooled, totalling approximately 3 mL. The sample was loaded on a HiLoad 16/600 Superdex 200 pg column pre-equilibrated with buffer M (500 mM KCl, 500 mM NaCl, 10 mM MgCl_2_, 1 mM β-mercaptoethanol, 0.002% mellitic acid, 50 mM Tris:HCl pH 8.0). The fractions containing His_6_-TEV-MESH1 were pooled and subjected to buffer exchange by repeated filtration with an Amicon Ultra (Millipore) centrifugal filter device (cut-off 15 kDa) pre-equilibrated in buffer N (100 mM NaCl, 5 mM MgCl_2_, 10% glycerol, 25 mM HEPES:KOH pH 7.5). The His-tag was cleaved off by adding TEV protease in a 1:100 molar ratio and the reaction mixture was incubated at 10 °C for overnight. After the His6 tag was cleaved off, the protein was passed though 1 mL HisTrap HP pre-equilibrated with buffer N. Fractions containing MESH1 in the flow-through were collected and concentrated on Amicon Ultra (Millipore) centrifugal filter device with 15 kDa cut-off (final concentration is 4.75 µM). The purity of protein preparations was assessed by SDS-PAGE and spectrophotometrically (OD_260_/OD_280_ ratio below 0.5). Protein preparations were aliquoted, frozen in liquid nitrogen and stored at –80 °C.

*C. marina* ATfaRel (VHp313, pET24d-His6-aTfaRel) was overexpressed in freshly transformed *E. coli* BL21 DE3. Fresh transformants were inoculated to final OD_600_ of 0.05 in the LB medium (800 mL) supplemented with 100 µg/mL kanamycin. The cultures were grown at 37 °C until an OD_600_ of 0.5, induced with 0.4 mM IPTG (final concentration) and grown for an additional one hour at 30 °C. The cells were harvested by centrifugation and resuspended in 30 ml of binding buffer (buffer O; 2 M NaCl, 5 mM MgCl_2_, 70 µM MnCl_2_, 50 mM arginine, 50 mM glutamic acid, 1 mM Mellitic acid, 20 mM imidazole, 10% glycerol, 4 mM β-mercaptoethanol, 25 mM HEPES:KOH pH 7.6) supplemented with 0.1 mM PMSF and 1 U/mL of DNase I, and cells were lysed by one passage through a high-pressure cell disrupter (Stansted Fluid Power, 150 MPa), cell debris was removed by centrifugation (35,000 rpm for 45 min, Type 45 Ti Beckman Coulter rotor) and clarified lysate was taken for protein purification.

Clarified cell lysate was filtered through a 0.2 µm syringe filter and loaded onto the HisTrap 1 mL HP column pre-equilibrated in buffer O (2 M NaCl, 5 mM MgCl_2_, 70 µM MnCl_2_, 50 mM arginine, 50 mM glutamic acid, 1 mM Mellitic acid, 20 mM imidazole, 10% glycerol, 4 mM β-mercaptoethanol, 25 mM HEPES:KOH pH 7.6). The column was washed with 5 CV of buffer O, and the protein was eluted with a linear gradient (10 CV, 0-100% buffer P) of buffer P (2 M NaCl, 5 mM MgCl_2_, 70 µM MnCl_2_, 50 mM arginine, 50 mM glutamic acid, 1 mM Mellitic acid, 500 mM imidazole, 4 mM β-mercaptoethanol, 25 mM HEPES:KOH pH 7.6). The fractions containing His_6_-ATfaRel were pooled and subjected to buffer exchange by repeated filtration with an Amicon Ultra (Millipore) centrifugal filter device (cut-off 3 kDa) pre-equilibrated in buffer Q (500 mM KCl, 5 mM MgCl_2_, 50 mM arginine, 50 mM glutamic acid, 10% glycerol, 4 mM β-mercaptoethanol, 25 mM HEPES:KOH pH 8.0). The purity of protein preparations was assessed by SDS-PAGE. Protein preparations were aliquoted, frozen in liquid nitrogen and stored at –80 °C.

## Preparation of *E. coli* fMet-tRNA_i_ ^fMet^ and tRNA^Phe^ modified by *Coprobacillus* sp. D7 FaRel2

fMet-tRNA_**i**_ ^fMet^ was prepared as described in before (Murina et al., 2018) using non-radioactive methionine.

To modify tRNA ^Phe^, the reaction mixture containing 5 µM tRNA^Phe^, 500 µM ATP and 50 nM FaRel2-FLAG_3_ in HEPES:Polymix buffer, pH 7.5 (Takada et al., 2020) (5 mM Mg^2+^ final concentration) supplemented with 1 mM DTT was incubated at 37 °C for 15 min. After that the reaction was supplemented with 0.1 volume of 3 M NaOAc (pH 4.6), the proteins were extracted with an equal volume of phenol/chloroform/isoamylalcohol (25:24:1), followed by a similar treatment with an equal volume of chloroform. As a negative control the same experiment was performed in the absence of FaRel2-FLAG_3_. The extracted tRNA was mixed with 2.5 volume of 95 % ethanol, precipitated at –20 °C overnight and pelleted by centrifugation. The pellet was washed with 100 µL of ice-cold 70% ethanol, dried at room temperature for 5 minutes, and dissolved in 5 mM KOAc (pH 5.1). The concentration of the purified tRNA was calculated by measuring the absorbance at 260 nm, and phosphorylation was validated by aminoacylation reaction (see below).

### Biochemical assays

*Cell-free translation:* experiments with PURExpress In Vitro Protein Synthesis Kit (NEB, E6800) were performed as per the manufacturer’s instructions with the addition of 0.8 U/µL RNase Inhibitor Murine (NEB, M0314S). FaRel2-FLAG_3_ was used at a final concentration of 50 nM, PhRel2-FLAG_3_ at 100 nM and His_6_-TEV-ATfaRel2 at 500 nM. As a control we used either HEPES:Polymix buffer, pH 7.5 (Takada et al., 2020) or eluate prepared from *E. coli* transformed with pBAD33 vector (mock). The total reaction volume was 6 µL per reaction for most of the experiments. To titrate the concentration of total tRNA in the reaction we used a combination of PURExpress Δ(aa, tRNA) Kit (NEB, E6840S) and total deacylated tRNA from *E. coli* MRE600 (Sigma-Aldrich, 10109541001). After incubation at 37 °C for the indicated time, the reaction mixture was mixed with 9-fold volume of 2x sample buffer (100 mM Tris:HCl pH 6.8, 4% SDS, 0.02% bromophenol blue, 20% glycerol, 20 mM DTT and 4% β-mercaptoethanol), and 5 µL of the mixture was resolved on 18% SDS-PAGE gel. The SDS-PAGE gel was fixed by incubating for 5 min at room temperature in 50% ethanol solution supplemented with 2% phosphoric acid, then stained and detected as mentioned in protein expression and purification.

*tRNA and oligonucleotide pyrophosphorylation by FaRel2*: the reaction conditions are described above, see ‘*Preparation of E. coli fMet-tRNA* ^*fMet*^ *and tRNA*^*Phe*^ *modified by* Coprobacillus *sp. D7 FaRel2*’. PhRel2-FLAG_3_ was used at a final concentration of 50 nM. Experiments with 5′-CACCN-3′ oligonucleotides used 50 µM oligonucleotides; tRNA (Chemical Block Ltd.) was used at a final concentration of 5 µM. The total reaction volume was either 8 or 20 µL per reaction. The reactions were started by the addition of 500 µM γ^32^P-ATP and incubated at 37 °C for either 10 or 30 minutes. To calculate the ratio of phosphorylation, the reaction mixture was mixed in 10% trichloroacetic acid supplemented with 70 ng/µL *E. coli* total tRNA as co-precipitant, kept on ice for 30 minutes, and centrifuged at 21,000 g for 30 minutes at 4 °C. After washing the pellet with 200 µL 10% TCA, tRNA was dissolved in 1 M Tris:HCl (pH 8.0) with shaking at 1,500 rpm for 20 min at 4 °C. The radioactivity was quantified by scintillation counting in 5 mL of EcoLite Liquid Scintillation Cocktail (MP Biomedicals).

To visualise phosphorylated tRNA the reaction sample was mixed in 2 volumes of RNA dye (98% formamide, 10 mM EDTA, 0.3% bromophenol blue and 0.3% xylene cyanol), tRNA was denatured at 37 °C for 10 min and resolved on urea-PAGE in 1 x TBE (8 M urea, 8% PAGE). The gel was stained with SYBR Gold (Life technologies, S11494) and exposed to an imaging plate overnight. The imaging plate was imaged by a Typhoon FLA 9500 (GE Healthcare). To visualise phosphorylated oligonucleotides the sample was resolved on urea-PAGE in 1 x TBE (5.6 M urea, 24% PAGE).

*Effects of pyrophosphorylation by FaRel2 on tRNA aminoacylation:* to probe the effect of FaRel2 on aminoacylation, 5 µM *E. coli* tRNA^Phe^ (Chemical Block Ltd.) was pre-incubated at 37 °C with or without 50 nM FaRel2 as well as 500 µM ATP for 10 minutes in HEPES:Polymix buffer, pH 7.5 (Takada et al., 2020) (5 mM Mg^2+^ final concentration) supplemented with 1 mM DTT and 160 µM ^3^H-phenylalanine. The total reaction volume was 20 µL. The tRNA was then aminoacylated by adding the same volume of aminoacylation mixture (4 mM ATP and 2 µM PheRS in HEPES:Polymix buffer (5 mM Mg^2+^ final concentration), 1 mM DTT) followed by additional incubation at 37 °C for 10 minutes. The reaction was quenched by adding trichloroacetic acid (TCA) to a final concentration of 10% as well as adding 70 ng/µL *E. coli* total tRNA as co-precipitant. After 30-minute incubation on ice, the tRNA was pelleted by centrifugation (21,000 g for 30 minutes at 4 °C). The supernatant was discarded, the pellet washed with 10% TCA and the tRNA was dissolved in 1 M Tris:HCl (pH 8.0) with shaking at 1,500 rpm for 20 min at 4 °C. Finally, ^3^H-radioactivity was quantified by scintillation counting in 5 mL of EcoLite Liquid Scintillation Cocktail (MP Biomedicals).

In the case of experiments using SAH enzymes, 5 µM tRNA^Phe^ with or without modification by FaRel2-FLAG_3_ was pre-incubated for 10 minutes with or without 1 µM MESH1, His_6_-ATfaRel or His_6_-TEV-ATfaRel2 at 37 °C in HEPES:Polymix buffer pH 7.5 (5 mM Mg^2+^ final concentration) (Takada et al., 2020) additionally supplemented with 1 mM MnCl_2_, 160 µM ^3^H-phenylalanine and 1 mM DTT. After 10 minute PheRS was added to final concentration of 1 µM and after additional 10 minutes at 37 °C the reaction was quenched by TCA and incorporation of the ^3^H-radioactivity was quantified by scintillation counting (see above).

^*3*^*H-ppGpp synthesis by* E. coli *RelA*: *E. coli* RelA synthase activity assay was performed in HEPES:Polymix buffer (5 mM Mg^2+^ final concentration) as described earlier (Takada et al., 2020) with minor modifications. Specifically, the assay was performed in the presence of 300 µM ^3^H-GDP, 1 mM ATP, 100 µM pppGpp, 50 nM native RelA and 100 nM 70S IC(MVF), in the presence or absence of 100 nM *E. coli* deacylated tRNA^Val^ (Chemical Block Ltd.); 5 µL total reaction volume per timepoint. *E. coli* deacylated tRNA^Val^ and ATP were initially incubated with either HEPES:Polymix buffer, empty vector lysate, or lysate containing FaRel2, at 37 °C for 30 minutes. GDP, pppGpp, and 70S IC were then added and the reactions were started by the addition of RelA. Subsequently, aliquots were withdrawn from the reaction mix, resolved on PEI-TLC, radioactivity quantified by scintillation counting and the turnovers were determined as previously described (Turnbull et al., 2019).

### Structural modelling and docking

The structure of FaRel2 was modelled using Rosetta (Song et al., 2013) based on the coordinates of *S. aureus* RelP (Manav et al., 2018) and *B. subtilis* RelQ (Steinchen et al., 2015). The models with the best scores were used for molecular docking as implemented in the web server version of HADDOCK (High ambiguity driven biomolecular docking) (van Zundert et al., 2016) together with the coordinates of yeast tRNA^Phe^ (Nissen et al., 1995).

For the docking procedure we defined residues in the active site; Y128 and the catalytic glutamine were selected as active residues (i.e. directly involved in the interaction). We then allowed HADDOCK to automatically select passive residues around the active residues. The program was run with default settings. The best cluster resulting from the docking experiment was selected to further probe the catalytic mechanism of FaRel2.

### Figure preparation

Figures were prepared using UCSF ChimeraX (Goddard et al., 2018), Igor Pro (WaveMetrics, Inc.), Adobe Illustrator (Adobe Inc.) and Adobe Photoshop (Adobe Inc.).

## QUANTIFICATION AND STATISTICAL ANALYSIS

Statistical analysis of tRNA aminoacylation, tRNA pyrophosphorylation and ^3^H-ppGpp synthesis data was performed using Igor Pro (WaveMetrics, Inc.). The data was plotted as individual data points as well as mean values ± standard deviations.

## DATA AND SOFTWARE AVAILABILITY

The study does not make use of unpublished data or software.

## RESOURCES TABLE

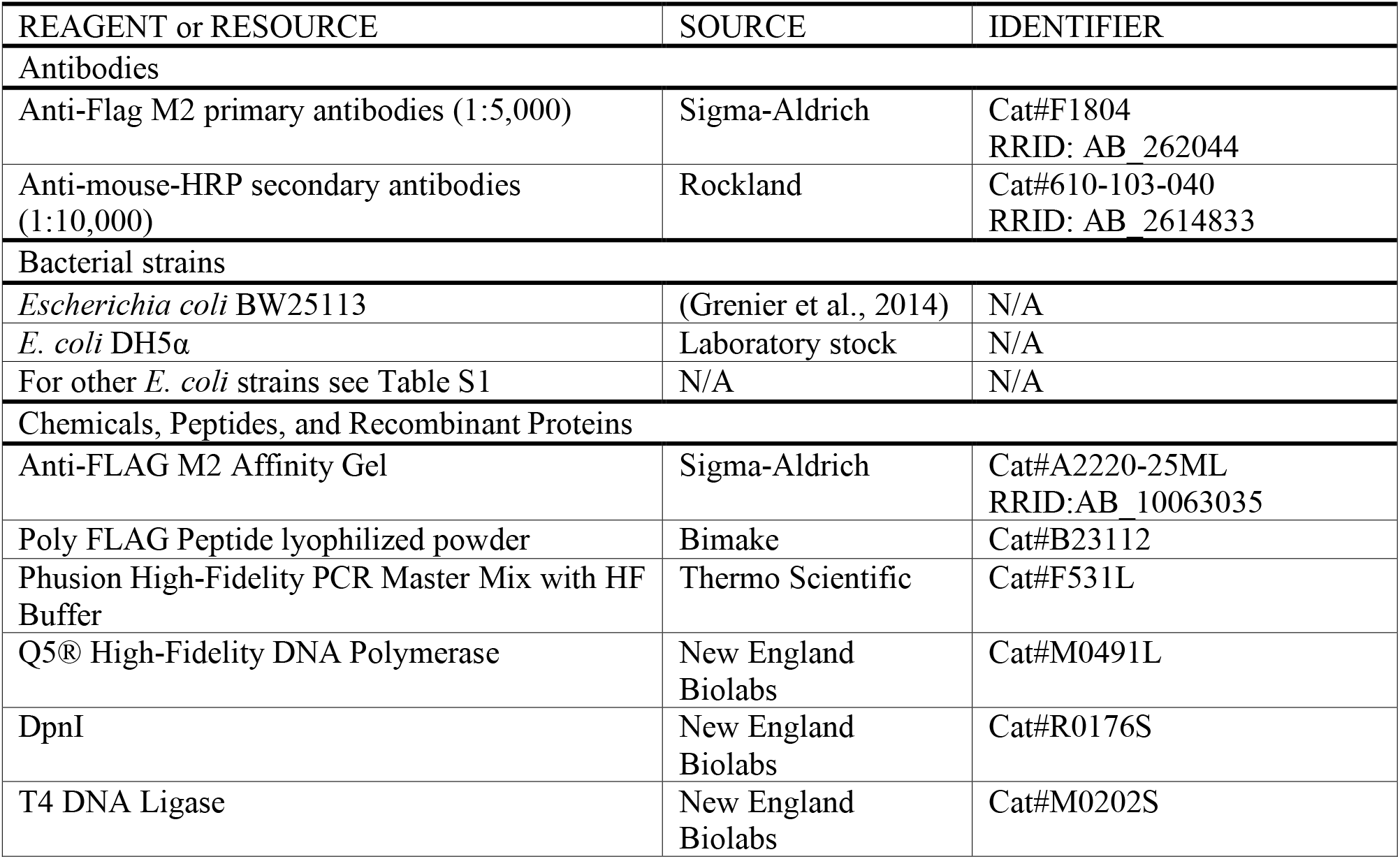
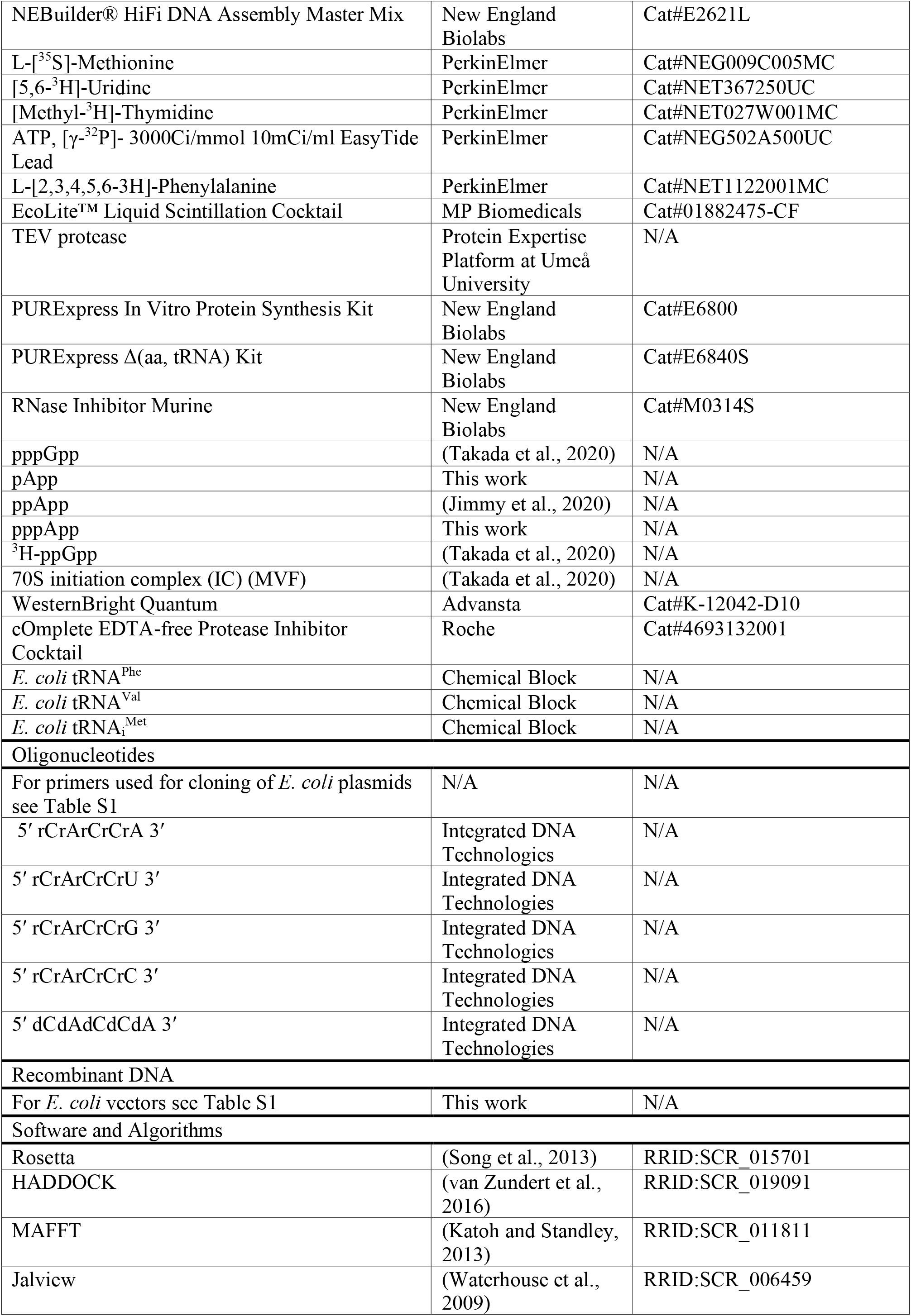
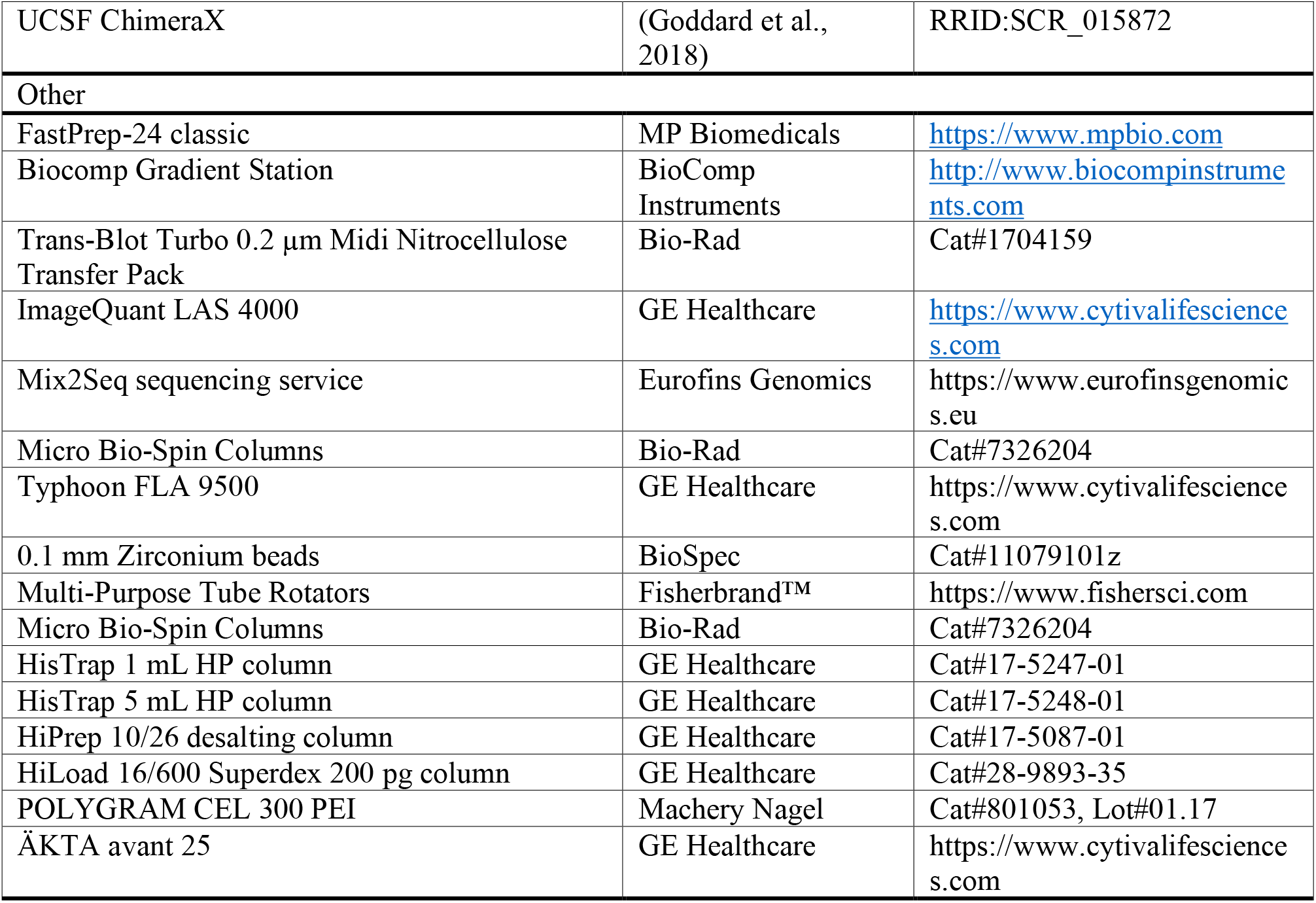

## DATA AND SOFTWARE AVAILABILITY

Sequences shown in the multiple sequence alignment of **Figure S5A** can be downloaded from the Uniprot database (https://www.uniprot.org/) in case of Rel and RelA, and the NCBI protein database for all other sequences, using the accession numbers shown in the figure.

## REFERENCES

Ahmad, S., Wang, B., Walker, M.D., Tran, H.R., Stogios, P.J., Savchenko, A., Grant, R.A., McArthur, A.G., Laub, M.T., and Whitney, J.C. (2019). An interbacterial toxin inhibits target cell growth by synthesizing (p)ppApp. Nature 575, 674–678.

Arenz, S., Abdelshahid, M., Sohmen, D., Payoe, R., Starosta, A.L., Berninghausen, O., Hauryliuk, V., Beckmann, R., and Wilson, D.N. (2016). The stringent factor RelA adopts an open conformation on the ribosome to stimulate ppGpp synthesis. Nucleic Acids Res 44, 6471–6481.

Atkinson, G.C., Tenson, T., and Hauryliuk, V. (2011). The RelA/SpoT homolog (RSH) superfamily: distribution and functional evolution of ppGpp synthetases and hydrolases across the tree of life. PLoS One 6, e23479.

Blower, T.R., Fineran, P.C., Johnson, M.J., Toth, I.K., Humphreys, D.P., and Salmond, G.P. (2009). Mutagenesis and functional characterization of the RNA and protein components of the toxIN abortive infection and toxin-antitoxin locus of Erwinia. J Bacteriol 191, 6029–6039.

Brosius, J., and Holy, A. (1984). Regulation of ribosomal RNA promoters with a synthetic lac operator. Proc Natl Acad Sci U S A 81, 6929–6933.

Brown, A., Fernandez, I.S., Gordiyenko, Y., and Ramakrishnan, V. (2016). Ribosome- dependent activation of stringent control. Nature 534, 277–280.

Burckhardt, R.M., and Escalante-Semerena, J.C. (2020). Small-Molecule Acetylation by GCN5-Related N-Acetyltransferases in Bacteria. Microbiol Mol Biol Rev 84.

Cai, Y., Usher, B., Gutierrez, C., Tolcan, A., Mansour, M., Fineran, P.C., Condon, C., Neyrolles, O., Genevaux, P., and Blower, T.R. (2020). A nucleotidyltransferase toxin inhibits growth of Mycobacterium tuberculosis through inactivation of tRNA acceptor stems. Sci Adv 6, eabb6651.

Candiano, G., Bruschi, M., Musante, L., Santucci, L., Ghiggeri, G.M., Carnemolla, B., Orecchia, P., Zardi, L., and Righetti, P.G. (2004). Blue silver: a very sensitive colloidal Coomassie G-250 staining for proteome analysis. Electrophoresis 25, 1327–1333.

Cashel, M., and Gallant, J. (1969). Two compounds implicated in the function of the RC gene of Escherichia coli. Nature 221, 838–841.

Castro-Roa, D., Garcia-Pino, A., De Gieter, S., van Nuland, N.A., Loris, R., and Zenkin, N. (2013). The Fic protein Doc uses an inverted substrate to phosphorylate and inactivate EF-Tu. Nat Chem Biol 9, 811–817.

Cheverton, A.M., Gollan, B., Przydacz, M., Wong, C.T., Mylona, A., Hare, S.A., and Helaine, S. (2016). A Salmonella Toxin Promotes Persister Formation through Acetylation of tRNA. Mol Cell 63, 86–96.

Cruz, J.W., Sharp, J.D., Hoffer, E.D., Maehigashi, T., Vvedenskaya, I.O., Konkimalla, A., Husson, R.N., Nickels, B.E., Dunham, C.M., and Woychik, N.A. (2015). Growth-regulating Mycobacterium tuberculosis VapC-mt4 toxin is an isoacceptor-specific tRNase. Nat Commun 6, 7480.

Dedrick, R.M., Jacobs-Sera, D., Bustamante, C.A., Garlena, R.A., Mavrich, T.N., Pope, W.H., Reyes, J.C., Russell, D.A., Adair, T., Alvey, R., et al. (2017). Prophage-mediated defence against viral attack and viral counter-defence. Nat Microbiol 2, 16251.

Ding, C.C., Rose, J., Sun, T., Wu, J., Chen, P.H., Lin, C.C., Yang, W.H., Chen, K.Y., Lee, H., Xu, E., et al. (2020). MESH1 is a cytosolic NADPH phosphatase that regulates ferroptosis. Nat Metab 2, 270–277.

Fiedoruk, K., Daniluk, T., Swiecicka, I., Sciepuk, M., and Leszczynska, K. (2015). Type II toxin-antitoxin systems are unevenly distributed among Escherichia coli phylogroups. Microbiology (Reading) 161, 158–167.

Fraikin, N., Goormaghtigh, F., and Van Melderen, L. (2020). Type II Toxin-Antitoxin Systems: Evolution and Revolutions. J Bacteriol 202.

Gaca, A.O., Colomer-Winter, C., and Lemos, J.A. (2015). Many means to a common end: the intricacies of (p)ppGpp metabolism and its control of bacterial homeostasis. J Bacteriol 197, 1146–1156.

Garcia-Pino, A., Zenkin, N., and Loris, R. (2014). The many faces of Fic: structural and functional aspects of Fic enzymes. Trends Biochem Sci 39, 121–129.

Geiger, T., Kastle, B., Gratani, F.L., Goerke, C., and Wolz, C. (2014). Two small (p)ppGpp synthases in Staphylococcus aureus mediate tolerance against cell envelope stress conditions. J Bacteriol 196, 894–902.

Goddard, T.D., Huang, C.C., Meng, E.C., Pettersen, E.F., Couch, G.S., Morris, J.H., and Ferrin, T.E. (2018). UCSF ChimeraX: Meeting modern challenges in visualization and analysis. Protein Sci 27, 14–25.

Goeders, N., Dreze, P.L., and Van Melderen, L. (2013). Relaxed cleavage specificity within the RelE toxin family. J Bacteriol 195, 2541–2549.

Grenier, F., Matteau, D., Baby, V., and Rodrigue, S. (2014). Complete Genome Sequence of Escherichia coli BW25113. Genome Announc 2.

Guzman, L.M., Belin, D., Carson, M.J., and Beckwith, J. (1995). Tight regulation, modulation, and high-level expression by vectors containing the arabinose PBAD promoter. J Bacteriol 177, 4121–4130.

Harms, A., Brodersen, D.E., Mitarai, N., and Gerdes, K. (2018). Toxins, Targets, and Triggers: An Overview of Toxin-Antitoxin Biology. Mol Cell 70, 768–784.

Harms, A., Stanger, F.V., Scheu, P.D., de Jong, I.G., Goepfert, A., Glatter, T., Gerdes, K., Schirmer, T., and Dehio, C. (2015). Adenylylation of Gyrase and Topo IV by FicT Toxins Disrupts Bacterial DNA Topology. Cell Rep 12, 1497–1507.

Haseltine, W.A., and Block, R. (1973). Synthesis of guanosine tetra- and pentaphosphate requires the presence of a codon-specific, uncharged transfer ribonucleic acid in the acceptor site of ribosomes. Proc Natl Acad Sci U S A 70, 1564–1568.

Hauryliuk, V., Atkinson, G.C., Murakami, K.S., Tenson, T., and Gerdes, K. (2015). Recent functional insights into the role of (p)ppGpp in bacterial physiology. Nat Rev Microbiol 13, 298–309.

Irving, S.E., Choudhury, N.R., and Corrigan, R.M. (2020). The stringent response and physiological roles of (pp)pGpp in bacteria. Nat Rev Microbiol.

Jaffe, A., Ogura, T., and Hiraga, S. (1985). Effects of the ccd function of the F plasmid on bacterial growth. J Bacteriol 163, 841–849.

Jaskolska, M., and Gerdes, K. (2015). CRP-dependent positive autoregulation and proteolytic degradation regulate competence activator Sxy of Escherichia coli. Mol Microbiol 95, 833–845.

Jimmy, S., Saha, C.K., Kurata, T., Stavropoulos, C., Oliveira, S.R.A., Koh, A., Cepauskas, A., Takada, H., Rejman, D., Tenson, T., et al. (2020). A widespread toxin-antitoxin system exploiting growth control via alarmone signaling. Proc Natl Acad Sci U S A 117, 10500–10510.

Jurenaite, M., Markuckas, A., and Suziedeliene, E. (2013). Identification and characterization of type II toxin-antitoxin systems in the opportunistic pathogen Acinetobacter baumannii. J Bacteriol 195, 3165–3172.

Jurenas, D., Chatterjee, S., Konijnenberg, A., Sobott, F., Droogmans, L., Garcia-Pino, A., and Van Melderen, L. (2017). AtaT blocks translation initiation by N-acetylation of the initiator tRNA(fMet). Nat Chem Biol 13, 640–646.

Katoh, K., and Standley, D.M. (2013). MAFFT multiple sequence alignment software version 7: improvements in performance and usability. Mol Biol Evol 30, 772–780.

Koonin, E.V., and Makarova, K.S. (2019). Origins and evolution of CRISPR-Cas systems. Philos Trans R Soc Lond B Biol Sci 374, 20180087.

Kudrin, P., Dzhygyr, I., Ishiguro, K., Beljantseva, J., Maksimova, E., Oliveira, S.R.A., Varik, V., Payoe, R., Konevega, A.L., Tenson, T., et al. (2018). The ribosomal A-site finger is crucial for binding and activation of the stringent factor RelA. Nucleic Acids Res 46, 1973–1983.

Kudrin, P., Varik, V., Oliveira, S.R., Beljantseva, J., Del Peso Santos, T., Dzhygyr, I., Rejman, D., Cava, F., Tenson, T., and Hauryliuk, V. (2017). Subinhibitory Concentrations of Bacteriostatic Antibiotics Induce relA-Dependent and relA-Independent Tolerance to beta-Lactams. Antimicrob Agents Chemother 61.

Lima-Mendez, G., Oliveira Alvarenga, D., Ross, K., Hallet, B., Van Melderen, L., Varani, A.M., and Chandler, M. (2020). Toxin-Antitoxin Gene Pairs Found in Tn3 Family Transposons Appear To Be an Integral Part of the Transposition Module. mBio 11.

Liu, K., Bittner, A.N., and Wang, J.D. (2015). Diversity in (p)ppGpp metabolism and effectors. Curr Opin Microbiol 24, 72–79.

Loveland, A.B., Bah, E., Madireddy, R., Zhang, Y., Brilot, A.F., Grigorieff, N., and Korostelev, A.A. (2016). Ribosome*RelA structures reveal the mechanism of stringent response activation. Elife 5.

Manav, M.C., Beljantseva, J., Bojer, M.S., Tenson, T., Ingmer, H., Hauryliuk, V., and Brodersen, D.E. (2018). Structural basis for (p)ppGpp synthesis by the Staphylococcus aureus small alarmone synthetase RelP. J Biol Chem 293, 3254–3264.

Murina, V., Kasari, M., Hauryliuk, V., and Atkinson, G.C. (2018). Antibiotic resistance ABCF proteins reset the peptidyl transferase centre of the ribosome to counter translational arrest. Nucleic Acids Res 46, 3753–3763.

Nanamiya, H., Kasai, K., Nozawa, A., Yun, C.S., Narisawa, T., Murakami, K., Natori, Y., Kawamura, F., and Tozawa, Y. (2008). Identification and functional analysis of novel (p)ppGpp synthetase genes in Bacillus subtilis. Mol Microbiol 67, 291–304.

Neidhardt, F.C., Bloch, P.L., and Smith, D.F. (1974). Culture medium for enterobacteria. J Bacteriol 119, 736–747.

Nissen, P., Kjeldgaard, M., Thirup, S., Polekhina, G., Reshetnikova, L., Clark, B.F., and Nyborg, J. (1995). Crystal structure of the ternary complex of Phe-tRNAPhe, EF-Tu, and a GTP analog. Science 270, 1464–1472.

Page, R., and Peti, W. (2016). Toxin-antitoxin systems in bacterial growth arrest and persistence. Nat Chem Biol 12, 208–214.

Patil, P.R., Vithani, N., Singh, V., Kumar, A., and Prakash, B. (2020). A revised mechanism for (p)ppGpp synthesis by Rel proteins: The critical role of the 2’-OH of GTP. J Biol Chem 295, 12851–12867.

Schattenkerk, C., Wreesmann, C.T., van der Marel, G.A., and van Boom, J.H. (1985). Synthesis of riboguanosine pentaphosphate ppprGpp (Magic Spot II) via a phosphotriester approach. Nucleic Acids Res 13, 3635–3649.

Schneider, C.A., Rasband, W.S., and Eliceiri, K.W. (2012). NIH Image to ImageJ: 25 years of image analysis. Nat Methods 9, 671–675.

Schureck, M.A., Dunkle, J.A., Maehigashi, T., Miles, S.J., and Dunham, C.M. (2015). Defining the mRNA recognition signature of a bacterial toxin protein. Proc Natl Acad Sci U S A 112, 13862–13867.

Senissar, M., Manav, M.C., and Brodersen, D.E. (2017). Structural conservation of the PIN domain active site across all domains of life. Protein Sci 26, 1474–1492.

Shimizu, Y., Inoue, A., Tomari, Y., Suzuki, T., Yokogawa, T., Nishikawa, K., and Ueda, T. (2001). Cell-free translation reconstituted with purified components. Nat Biotechnol 19, 751–755.

Song, S., and Wood, T.K. (2020). A Primary Physiological Role of Toxin/Antitoxin Systems Is Phage Inhibition. Front Microbiol 11, 1895.

Song, Y., DiMaio, F., Wang, R.Y., Kim, D., Miles, C., Brunette, T., Thompson, J., and Baker, D. (2013). High-resolution comparative modeling with RosettaCM. Structure 21, 1735–1742.

Sprinzl, M., and Richter, D. (1976). Free 3’-OH group of the terminal adenosine of the tRNA molecule is essential for the synthesis in vitro of guanosine tetraphosphate and pentaphosphate in a ribosomal system from Escherichia coli. Eur J Biochem 71, 171–176.

Steinchen, W., Schuhmacher, J.S., Altegoer, F., Fage, C.D., Srinivasan, V., Linne, U., Marahiel, M.A., and Bange, G. (2015). Catalytic mechanism and allosteric regulation of an oligomeric (p)ppGpp synthetase by an alarmone. Proc Natl Acad Sci U S A 112, 13348–13353.

Steinchen, W., Vogt, M.S., Altegoer, F., Giammarinaro, P.I., Horvatek, P., Wolz, C., and Bange, G. (2018). Structural and mechanistic divergence of the small (p)ppGpp synthetases RelP and RelQ. Sci Rep 8, 2195.

Sun, D., Lee, G., Lee, J.H., Kim, H.Y., Rhee, H.W., Park, S.Y., Kim, K.J., Kim, Y., Kim, B.Y., Hong, J.I., et al. (2010). A metazoan ortholog of SpoT hydrolyzes ppGpp and functions in starvation responses. Nat Struct Mol Biol 17, 1188–1194.

Takada, H., Roghanian, M., Murina, V., Dzhygyr, I., Murayama, R., Akanuma, G., Atkinson, G.C., Garcia-Pino, A., and Hauryliuk, V. (2020). The C-terminal RRM/ACT domain is crucial for fine-tuning the activation of ‘long’ RelA-SpoT Homolog enzymes by ribosomal complexes. Frontiers in microbiology 11, 277.

Turnbull, K.J., Dzhygyr, I., Lindemose, S., Hauryliuk, V., and Roghanian, M. (2019). Intramolecular Interactions Dominate the Autoregulation of Escherichia coli Stringent Factor RelA. Front Microbiol 10, 1966.

van Zundert, G.C.P., Rodrigues, J., Trellet, M., Schmitz, C., Kastritis, P.L., Karaca, E., Melquiond, A.S.J., van Dijk, M., de Vries, S.J., and Bonvin, A. (2016). The HADDOCK2.2 Web Server: User-Friendly Integrative Modeling of Biomolecular Complexes. J Mol Biol 428, 720–725.

Varik, V., Oliveira, S.R.A., Hauryliuk, V., and Tenson, T. (2017). HPLC-based quantification of bacterial housekeeping nucleotides and alarmone messengers ppGpp and pppGpp. Sci Rep 7, 11022.

Waterhouse, A.M., Procter, J.B., Martin, D.M., Clamp, M., and Barton, G.J. (2009). Jalview Version 2--a multiple sequence alignment editor and analysis workbench. Bioinformatics 25, 1189–1191.

Winther, K.S., and Gerdes, K. (2011). Enteric virulence associated protein VapC inhibits translation by cleavage of initiator tRNA. Proc Natl Acad Sci U S A 108, 7403–7407.

Xiao, H., Kalman, M., Ikehara, K., Zemel, S., Glaser, G., and Cashel, M. (1991). Residual guanosine 3’,5’-bispyrophosphate synthetic activity of relA null mutants can be eliminated by spoT null mutations. J Biol Chem 266, 5980–5990.

Yamaguchi, Y., and Inouye, M. (2009). mRNA interferases, sequence-specific endoribonucleases from the toxin-antitoxin systems. Prog Mol Biol Transl Sci 85, 467–500.

Zhu, M., Pan, Y., and Dai, X. (2019). (p)ppGpp: the magic governor of bacterial growth economy. Curr Genet 65, 1121–1125.

